# Rapid switching of in-phase and anti-phase interhemispheric rhythms within a single theta cycle: a mechanistic explanation

**DOI:** 10.64898/2026.10.23.761406

**Authors:** Sameer Kailasa, Izabela Jedrasiak-Cape, Daniele Avitabile, Victoria Booth, Omar J. Ahmed

## Abstract

The granular retrosplenial cortex (RSG) supports memory consolidation and spatial navigation. *In vivo* recordings of local field potential (LFP) from RSG reveal coexisting gamma (30-80 Hz) and spline (100-160 Hz) oscillations, coupled to distinct phases of the background theta (∼8 Hz) rhythm. Remarkably, bilateral recordings reveal that gamma and splines exhibit divergent interhemispheric phase relationships: gamma is in-phase across hemispheres, while splines are anti-phase. Here, using numerical bifurcation analysis of next-generation neural mass models, we show that RSG oscillations within a single hemisphere can be understood as alternation of PING (pyramidal-interneuron gamma) and ING (interneuron gamma) mechanisms in an excitatory-inhibitory network. Theta-frequency external drive to the inhibitory population switches the network between these two rhythms. Next, we build canonical interhemispheric circuits, composed of two such excitatory-inhibitory modules, delay-coupled across hemispheres. These circuits switch between in-phase gamma and anti-phase splines within a single theta cycle, reproducing the cross-spectral statistics of experimental RSG recordings. Finally, we present analytical results for threshold-linear neural mass models, showing cross-hemispheric phase patterning is governed by the ratio of cross-hemispheric delay to local inhibitory delay; we obtain explicit ranges of this ratio for which RSG-like patterning could be expected. Our work establishes that the unique oscillatory profile of RSG can be explained parsimoniously by minimal interhemispheric circuit configurations and provides a framework for experimental interrogation of interhemispheric rhythms.

**Significance statement:** The granular retrosplenial cortex (RSG) contributes to our sense of spatial orientation. Recordings from RSG reveal nested rhythmic activity, with two fast rhythms, gamma and splines, embedded within a slower theta rhythm. Intriguingly, gamma rhythms are synchronized across hemispheres (in-phase), while splines are antisynchronized (anti-phase): spline peaks in one hemisphere correspond with troughs in the other. Here, we present and analyze mathematical models which capture how the observed oscillation patterns can arise out of simple neural circuit configurations. These models make testable predictions about what network architectures could produce the oscillations, and how the oscillations are controlled by external inputs to RSG, supporting future experiments which aim to better understand left/right brain communication in health and disease.

## Introduction

The granular retrosplenial cortex (RSG) is an important node of the neural circuitry supporting spatial navigation and memory ([1], [2], [3], [4], [5]). A key member of the densely connected default mode network, its role integrating hippocampal outputs (via dorsal subiculum; [6], [7]) with subcortical outputs (via anterior thalamus; [8], [9]) positions RSG as a key player in the tripartite interactions which subserve episodic memory ([9]). Macroscopic oscillations observable in RSG field potentials ([10], [11], [12]) provide intriguing signatures of the underlying thalamo-cortico-hippocampal dynamics.

During active exploration behavior and REM sleep, extracellular voltage recordings from superficial layers of RSG exhibit a prominent theta rhythm (4-12 Hz) and superimposed oscillations in the gamma (30-80 Hz, also called slow gamma) and spline/HFO (100-160 Hz high frequency oscillations, also called fast gamma) frequency bands. Both gamma and splines have amplitudes modulated by the background theta rhythm, with gamma preferentially occurring during the rising phase of theta, and splines preferentially occurring at the peak and falling phase of RSG theta ([10], [11], [12]). Remarkably, gamma and splines display divergent cross-hemispheric phase relationships: bilateral local field potential (LFP) recordings reveal that gamma oscillations are in-phase across hemispheres, while splines are anti-phase ([10]). Accordingly, splines were named in analogy with mechanical splines – the anti-phase teeth on interlocking gears ([10]).

A panoply of neuroscience literature, both experimental and theoretical, indicates the phenomenological ubiquity and computational significance of long-range oscillatory synchronization, especially in the gamma band ([13], [14], [15], [16]). Indeed, oscillations have been implicated in feature binding ([17], [18]), attentional allocation ([19]), and memory consolidation ([20], [21]). Thus, with the goal of ultimately characterizing the functions performed by a particular neural circuit, a natural step is to first establish how its macroscopic oscillations might constrain or support these functions. For RSG, this amounts to asking: what kind of brain circuits might produce the oscillatory patterning observed in the LFPs?

Here, we explore the hypothesis, consistent with prior experimental findings in subiculum ([22]), that RSG gamma and spline oscillations are respectively generated by PING-(pyramidal-interneuron gamma, supported by an excitatory → inhibitory → excitatory feedback loop; [23]) and ING-like (interneuron gamma, supported by an inhibitory → inhibitory feedback loop; [24], [25]) mechanisms. In networks capable of producing both PING and ING oscillations, variations in the external inputs can mediate transitions between regimes dominated by either oscillation type; this phenomenon is called *PING-ING alternation* ([26], [27]). Accordingly, one possibility is that theta-gamma phase-amplitude coupling in RSG arises via alternation of locally generated PING and ING rhythms, driven by theta-rhythmic inhibition of RSG interneurons from medial septum ([8]).

Subject to this PING-ING alternation hypothesis, the RSG recordings impose an interesting additional constraint on the dynamics: PING/gamma oscillations are in-phase across hemispheres, while ING/spline oscillations are anti-phase across hemispheres. Accordingly, in this work, we seek to identify minimal interhemispheric network configurations which can co-express in-phase PING and anti-phase ING oscillations, and study their dynamics under theta-rhythmic periodic forcing. Our approach utilizes the established mean-field framework afforded by globally coupled quadratic integrate-and-fire (QIF) neurons ([28], [25], [29], [30], [31], [32]). Importantly, the dynamics of these QIF networks are captured exactly by a set of low-dimensional mean field equations, which can then be probed directly using techniques from bifurcation theory.

Herein, we use the mean-field framework to successfully construct model networks capable of PING-ING alternation, exhibiting divergent phase relationships across hemispheres, as observed *in vivo*. In fact, we find that either delayed cross-hemispheric E → I or I → I projections can suffice to qualitatively recapitulate the experimentally observed patterning, but that E → I projections reproduce a separation of PING and ING frequencies more consistent with experimental observations. Moreover, we show that a key quantity determining the cross-hemispheric phase relationship for PING and ING oscillations simultaneously is the ratio of cross-hemispheric delay to the within-hemisphere inhibitory feedback delay.

A key experimental challenge in neuroscience is to characterize the macroscopic dynamical regimes occupied by specific brain networks, since these regimes strongly constrain intra- and inter-regional communication. We anticipate our study will inform the design and interpretation of future experiments aimed at probing interhemispheric brain communication in health and disease, in RSG and beyond.

## Materials and Methods

In this section, we first introduce globally-coupled networks of QIF neurons with microscopic Cauchy white noise, along with the corresponding mean-field formulation (quadratic-integrate-and-fire neural mass; QIF-NM). We derive the transfer function of the mean-field network and use this to obtain defining equations for local bifurcations of a network steady state. Then, we specialize to the case of symmetric interhemispheric networks and describe the minimal interhemispheric networks of interest in this work. Finally, we describe sinusoidal theta forcing provided to the mean-field network, along with procedures for data analysis.

### Microscopic QIF network and mean-field reduction

We consider 1 ≤ *α* ≤ *P* populations of *quadratic integrate-and-fire* (QIF) neurons, with neurons in population *α* indexed by 1 ≤ *j* ≤ *N*_*α*_. Neurons within and across populations are globally coupled via chemical synapses. Specifically, the state of each neuron (*j, α*) is given by a dimensionless voltage variable *V*_*j,α*_ satisfying the stochastic differential equation

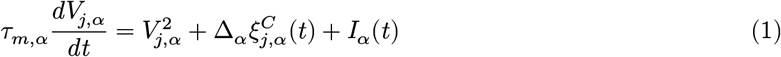

Here, *τ*_*m,α*_ is the population-specific membrane time constant; each *ξ*^*C*^ is a neuron-specific Cauchy white noise; Δ_*α*_ denotes the population-specific microscopic noise amplitude; and *I*_*α*_(*t*) is the total input current to neurons in population *α*. We follow the standard convention ([28], [30]) for a QIF neuron to *spike* when its voltage blows up to ∞ in finite time, and upon spike instantaneously reset the voltage to −∞. Relevant macroscopic observables associated with each population are the *population mean voltage*

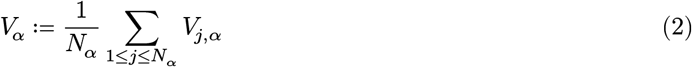

and the *population mean firing rate*

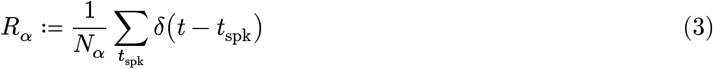

where the latter sum runs over all spikes emitted by members of *α*. The input current *I*_*α*_ is comprised by

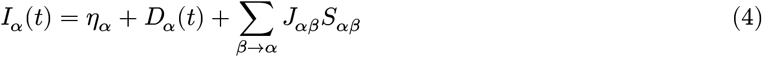

where *η*_*α*_ and *D*_*α*_(*t*) denote DC (time-constant) and AC (time-varying) components of an external forcing input. Meanwhile ∑_*β*→*α*_ *J*_*αβ*_*S*_*αβ*_ is the recurrent synaptic input from the network, with *J*_*αβ*_ denoting the directed chemical coupling strength to population *α*, from population *β*. Finally, in order to couple the populations to one another, we conclude the specification of the model by taking *Q*_*αβ*_*S*_*αβ*_ = π*τ*_*m,β*_*R*_*β*_ for synapse-specific linear-time-invariant operators *Q*_*αβ*_ which encode the synaptic kinetics ([33]).

In the thermodynamic limit *N*_*α*_ → ∞, the state of each population is described by a time-varying distribution *P*_*α*_(*v, t*) over voltages, satisfying a fractional Fokker-Planck equation ([34], [35]). Remarkably, in the presence of dissipation (Δ_*α*_ > 0 for all 1 ≤ *α* ≤ *P* ), the voltage distribution for each population asymptotically collapses onto the so-called *Ott-Antonsen* submanifold comprised of Cauchy distributions, such that

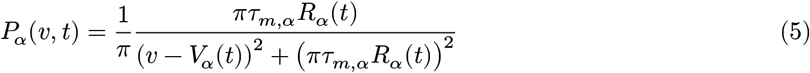

holds after an initial transient ([30], [34], [35]). Here *V*_*α*_ and *R*_*α*_ correspond precisely to limiting (as *N*_*α*_ → ∞) values of the population mean voltage and population mean firing rate from above. Substituting the Ott-Antonsen distributional form into the fractional Fokker-Planck equation, one derives macroscopic equations of evolution for *R*_*α*_ and *V*_*α*_, which take the form

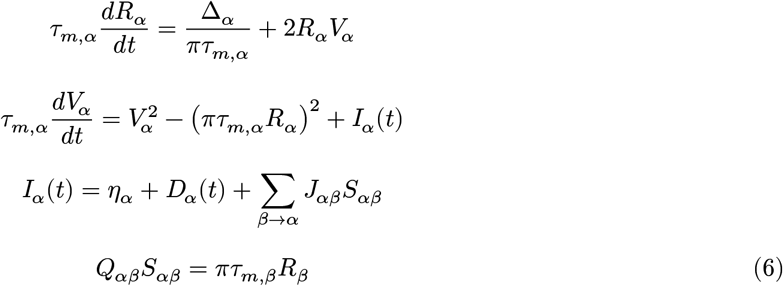

These mean-field dynamics are deterministic; to emulate stochastic finite-size effects, we make the Poissonian rate approximation and introduce population output stochasticity into the synaptic transfer as

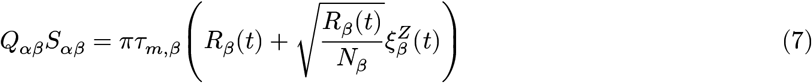

where 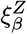 is a standard Gaussian white noise (independent across populations, and we interpret multiplicative noise in the Îto sense). Note that this approximation does neglect resonant peaks known to occur at low frequencies in the true population output shot noise ([36]); nonetheless, it still provides a reasonable mesoscopic network model ([37]). Lastly, noting that the microscopic noise amplitude Δ_*α*_ sets a voltage scale for each population, we rescale the parameters and variables to eliminate this degree of freedom:

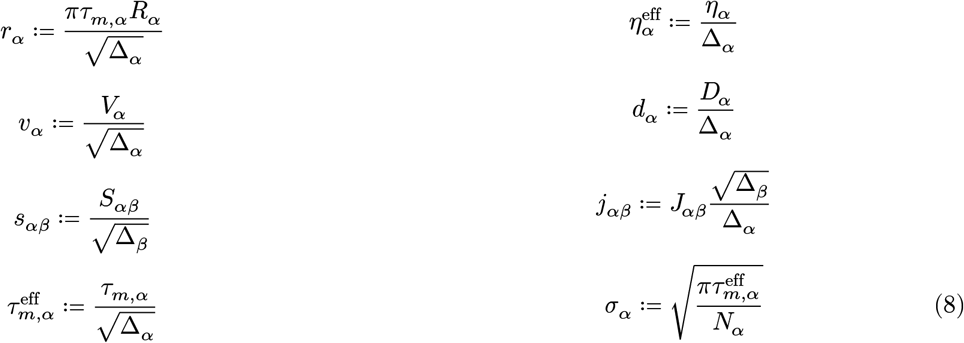

Equivalently, we can (and do) set Δ_*α*_ = 1 for all populations, without loss of generality. Thus, dropping the “eff” superscripts to lighten notation, we arrive at the final *QIF neural mass equations* (**QIF-NM**):

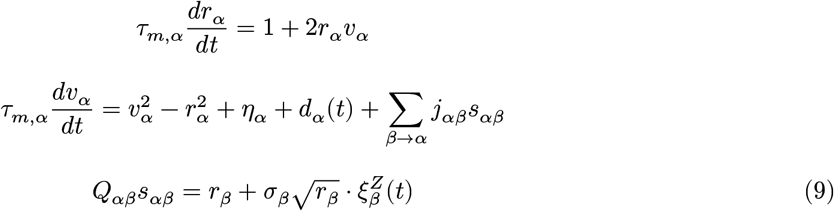

In absence of AC forcing (*d*_*α*_(*t*) = 0) and finite-size noise (*σ*_*α*_ = 0), we denote a steady state of Equation 9 by 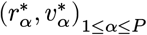, which satisfies the system

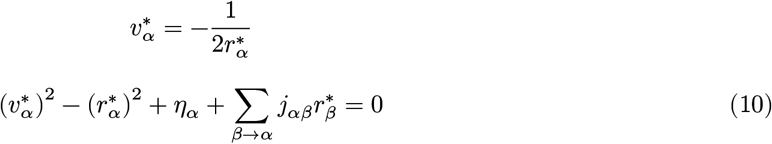

The steady state voltages are determined entirely by the rates, so the equilibrium can be equally well denoted 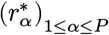.

### Network transfer function and synaptic kinetics

Let 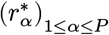 be a steady state of Equation 9, which we assume to be stable. To fix notation, let 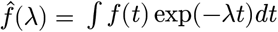 denote (formally) the two-sided Laplace transform of a signal *f*. We are interested in the evolution of a small perturbation (*δr*_*α*_, *δv*_*α*_)_1≤α≤*P*_ about the given steady state; this perturbation satisfies the linearized tangent dynamics

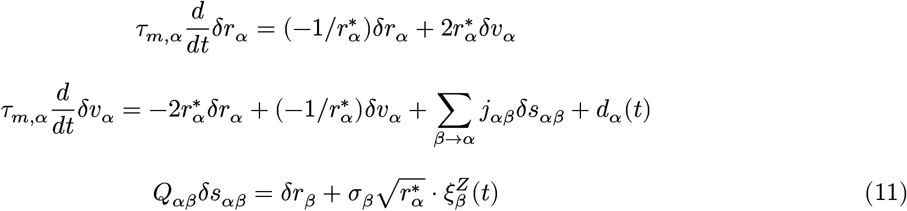

so long as the AC drives *d*_*α*_ and macroscopic noise amplitudes *σ*_*α*_ are small. Note that this linearization describes simply a network of spiral sink oscillators, each of whose parameters depend on the steady state negotiated via network couplings. Taking the Laplace transform, we write

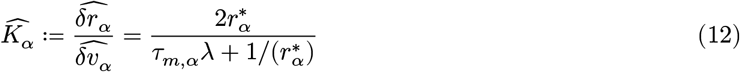

for the voltage-to-rate transfer function and

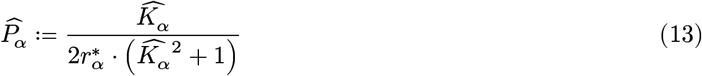

for the overall voltage response function of the uncoupled population. Denoting 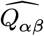 for the Laplace transform of the synaptic operator, we obtain

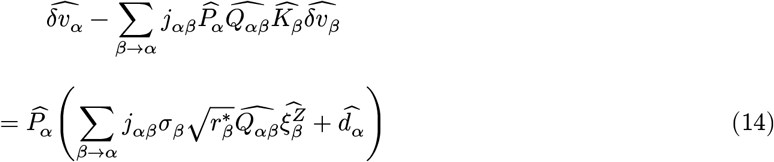

which reveals the distinct contributions of AC drive and finite-size noise to total network power. We set 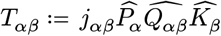 and 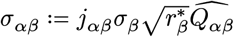, taking ***T*** = (*T*_αβ_ )_1≤*α,β*≤*P*_ and ***σ*** = (*σ*_αβ_ )_1≤*α,β*≤*P*_ to denote the matrices filled with these entries. Furthermore, let 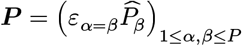 be the diagonal matrix which stacks the population response functions (*ε* here denoting an indicator function). Then the voltage fluctuations are

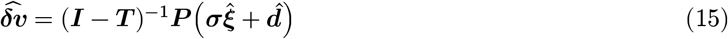

By setting *λ* = *iω*, we recover the Fourier transform, and hence the cross-spectral density of voltage fluctuations results as

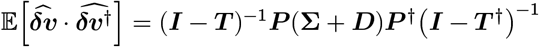

where ***Σ*** = ***σσ***^†^ is the cross-spectral density of the synaptically filtered finite-size noise and 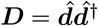 is the cross-spectral density of the external AC drive. Furthermore, local bifurcations of the steady state 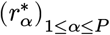 are obtained through the characteristic equation

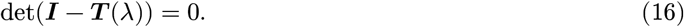

In particular, det(***I*** − ***T*** (0)) = 0 yields the locus of fold bifurcations, while det(***I*** − ***T*** (*iω*)) = 0 yields the Hopf locus. This calculation can be compared with equation (35) in [38], which carries out the analogous computation for networks of rate models and spiking networks of leaky integrate-and-fire neurons.

In this work, we assume the synaptic transfer 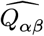 arises as a composition of first-order kinetics with a distributed synaptic delay. For simplicity, we assume the decay times of the first-order kinetics depend only on the presynaptic population, while the mean delay may depend on both pre- and postsynaptic populations. We assume Gamma distributed delays, such that we can utilize the linear chain trick; this trick allows to evaluate the infinite-dimensional synaptic filter induced by the Gamma delay distribution as an exact finite-dimensional linear filter. The models we construct are thereby systems of ordinary differential equations, and not delay differential equations, simplifying the numerical analyses ([39], [40]). More precisely, in the notation introduced above, we consider the synaptic response

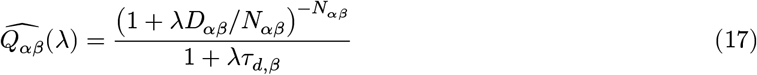

where *D*_*αβ*_ is the mean synaptic delay, *N*_*αβ*_ is the length of the approximating linear chain, and *τ*_*d,α*_ is the synaptic decay time. Note that:

- a chain length of *N*_*αβ*_ corresponds to a delay distribution with coefficient of variation 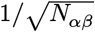,
- the numerator of 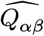 converges to the linear response for an exact delay exp(−*λD*_*αβ*_) as *N*_*αβ*_→ ∞,
- 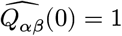, and thus delays will generally affect Hopf but not fold loci.

In [39], the authors show that the linear chain approximation to exact delays preserves the local bifurcation structure of the infinite-dimensional system, while possibly modifying the global bifurcation structure. Since distributed delays are certainly expected in the brain, we have elected to focus on this model formulation.

### Symmetric interhemispheric networks

In this work, we will focus on a minimal *symmetric interhemispheric network*, which we define as a network with 1≤ *α* ≤ 2*P* populations such that:

- population *α* has the same intrinsic parameters as population *α* + *P* for all 1 ≤ *α* ≤ *P*,
- the weight matrix takes the block form

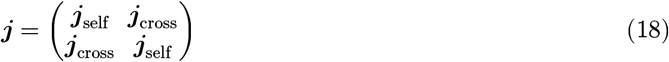

- the mean delay matrix ***D*** *≔* (*D*_*αβ*_)_1≤*α,β*≤*P*_ takes the block form

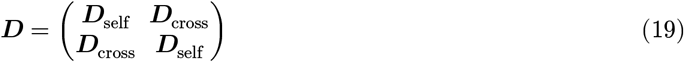

Subject to these assumptions, the network equations are invariant under the exchange of populations *α* and *α* + *P* for 1 ≤ *α* ≤ *P* . Implicitly, we interpret populations 1 ≤ *α* ≤ *P* as belonging to one hemisphere and *P* + 1 ≤ *α* ≤ 2*P* as belonging to the other.

We consider that the network admits a stable symmetric steady state, i.e. a state for which 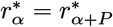 for 1 ≤ *α* ≤ *P* . Under the above assumptions, the network response matrix ***T*** about this steady state itself takes a block matrix form:

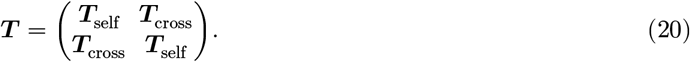

Using the identity

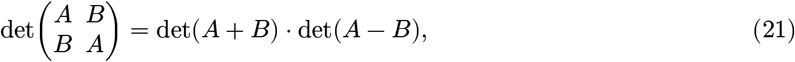

the characteristic equation det(***I*** − ***T*** (*λ*)) = 0 factors as

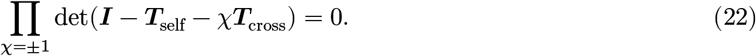

Solutions to det(***I*** − ***T***_self_ − ***T***_cross_) = 0 are called *longitudinal instabilities*, while solutions to det(***I*** − ***T***_self_ + ***T***_cross_) = 0 are called *transverse instabilities*. Note that longitudinal instabilities correspond precisely to those of a reduced single-hemisphere network, while transverse instabilities reflect states which break either temporal or spatial symmetry, and exist only in the full network.

In what follows, we make the simplifying assumption that ***D***_self_ = **0** and ***D***_cross_ = *D***1** where **0** and **1** denote the all zeros and all ones matrix, respectively. The key parameter *D* is the *mean interhemispheric delay*. Furthermore, note that symmetric interhemispheric networks admit a convenient re-parameterization in terms of the *total synaptic weight* of a connection type, and the *interhemispheric weight scaling* which quantifies the fraction of total weight which is assigned to the corresponding interhemispheric synapse (versus the corresponding local synapse). More precisely, we set ***j***_tot_ = ***j***_self_ + ***j***_cross_ and **0** ≤ ***ρ*** *≔* ***j***_cross_/***j***_tot_ ≤ **1** where the division and inequalities are interpreted element-wise. Observe that ***ρ*** is not well-defined for zero entries of ***j***_tot_. When ***ρ*** = **0**, the hemispheres are wholly decoupled, while when ***ρ*** = **1**, all synapses are cross-hemispheric. We emphasize that the steady state network firing rates (for a symmetric steady state) depend only on ***j***_tot_, such that the total weight matrix sets the overall activity level; meanwhile, the ***ρ*** matrix only redistributes this activity between local and cross-hemispheric pathways.

### Minimal network architectures coexpressing PING/ING

Within the QIF-NM framework introduced above, the minimal network capable of generating both PING and ING oscillations consists of a single excitatory-inhibitory pair of neural masses. The resulting mean-field equations are five-dimensional, with the synaptic decay time *τ*_*d,E*_ = 0 for excitatory outputs set to zero, while a nonzero inhibitory synapse time *τ*_*d,I*_ > 0 is necessary for the network to possess an ING instability ([25]). Accordingly, we obtain the **2POP** equations

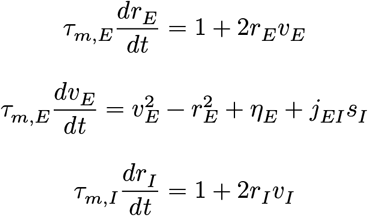

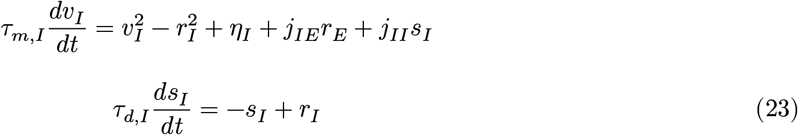

as our description of each hemisphere (representing, in the biological interpretation, activity of layer 2/3 pyramidal cells and fast-spiking interneurons of RSG). The assumption that *τ*_*d,E*_ ≪ *τ*_*d,I*_ is justified by slice measurements of synaptic kinetics in RSG ([41], [42]). Note that this model formulation excludes recurrent excitation. Although recurrent excitation is necessary to generate a PING instability in the classical 2D Wilson-Cowan ODE ([43]), it is not strictly required in the presence of inhibitory delays. We acknowledge many cortical networks do operate in an inhibition stabilized regime, where moderate recurrent excitation is balanced by strong feedback inhibition ([44]). Nonetheless, experiments have observed no recurrent excitation in pairs comprised of the dominant pyramidal cell type of superficial RSG ([41], [7], [42]), despite the apparent contribution of these layers to RSG high-frequency oscillations ([11]). Experiments also indicate the absence of monosynaptic cross-hemispheric excitation within this dominant pyramidal cell type ([7], [42]), such that overall the level of recurrent excitation in this circuit appears low. We focus in this paper on the limiting case *j*_*EE*_ = 0, noting that the **2POP** network admits a unique steady state for all *j*_*EE*_ sufficiently small (see Addendum 1) and that numerical experiments indicate similar dynamics throughout this parameter regime.

Our minimal interhemispheric circuit consists of four populations, an excitatory-inhibitory **2POP** pair in each hemisphere. Thus, we study the **4POP** equations

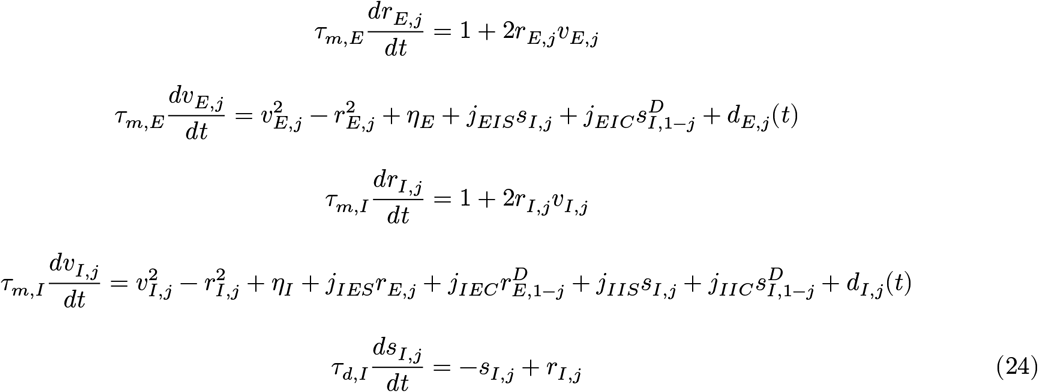

for *j* = 0, 1. Here *f* ^*D*^ for a signal *f*(*t*) denotes the linear-chain filtered version of *f*, i.e. *f* ^*D*^ *≔ f*_*N*_ where

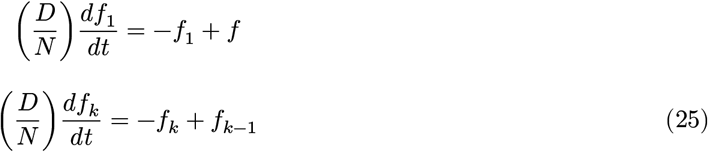

for 2 ≤ *k* ≤ *N* and *N* denotes the length of the approximating chain ([39]). This construction yields a finite-dimensional approximation to the time-delayed signal, i.e. *f* ^{*D*}^(*t*) ≈ *f*(*t* − *D*). For a linear chain of length *N*, the total **4POP** system has dimension 4*N* + 10. Here, subscripts *S* denote ipsilateral projections, while subscripts *C* denote contralateral projections. If we assume one interhemispheric projection type dominates (i.e. only one of *j*_*XY C*_ > 0 for (*X, Y* ) = (*E, I*), (*I, E*), or (*I, I*), similar to the approach of [45]), then this dimensionality reduces to 2*N* + 10. Here we limit the scope to the latter setup, with *N* = 5 (i.e. a Gamma delay distribution with 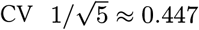, resulting in a 20-dimensional system of ordinary differential equations. By construction, the **4POP** system is a symmetric interhemispheric network, as defined above. Utilizing the alternative parameterization of **4POP**, we set *j*_*XY*_ = *j*_*XY S*_ + *j*_*XY C*_ and 0 ≤ *ρ*_*XY*_ = *j*_*XY C*_/*j*_*XY*_ ≤ 1 for all connection types *XY* . We refer to the **4POP** model for which only *ρ*_*XY*_ is nonzero simply as the **4POP-XY** model. For example, **4POP-IE** describes the network whose only cross-hemispheric projections are from ipsilateral excitatory cells to contralateral inhibitory cells.

A useful fact, which somewhat simplifies the study of these models, is that cross-hemispheric E → I projections and I → E projections admit precisely the same local bifurcations of the symmetric steady state (see Addendum 2). We emphasize that this does not preclude the possibility of distinct global dynamics between the **4POP-EI** and **4POP-IE** models, nor does it imply that Hopf points admit the same criticality across models. Nonetheless, one expects that in and close to the linear model regime, dynamics of these two cases are similar. Thus, in this work, we restrict our focus to comparing the **4POP-IE** and **4POP-II** models. All figures in this paper utilize the default parameter set *τ*_*m,E*_ = 9 ms, *τ*_*m,I*_ = 5 ms, *τ*_*d,I*_ = 1 ms, *j*_*EI*_ = −14, *j*_*IE*_ = 20, and *j*_*II*_ = −14. However, we vary the mean interhemispheric delay *D*, as well as the interhemispheric weight scaling *ρ*_*IE*_ and *ρ*_*II*_ .

### Form of external inputs, data analysis, and numerics

We assume the AC input to each population is a harmonic periodic forcing, namely *d*_*α*_(*t*) = *A*_*α*_ sin(2π*t*/*T*_*α*_), where *A*_*α*_ is the forcing amplitude and *T*_*α*_ is the forcing period. In this work, we aim to model symmetric theta-rhythmic external inputs from medial septum to RSG, which likely target interneurons preferentially ([8]). Thus, we set *A*_*E*,0_ = *A*_*E*,1_ = 0, *A*_*I*,0_ = *A*_*I*,1_ *≕ A*_*I*_, and *T*_*I*,1_ = *T*_*I*,2_ = 150 ms.

In order to quantify the oscillatory content of both experimental and model-generated data, we employ the continuous wavelet transform, as implemented in the Python package fCWT ([46]). In particular, we use the default complex Morlet wavelet with wavelet parameter *σ* = 2. Let CWT_*s*_(*ω, t*) denote the continuous wavelet transform of signal *s*, at frequency *ω* and time *t*. We utilize the wavelet cross-spectrum defined as

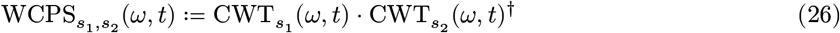

where *s*_1_, *s*_2_ are the two signals of interest and † denotes complex conjugation.

When analyzing experimental data, *s*_1_ and *s*_2_ are always two cross-hemispheric LFP channels. When analyzing model data, *s*_1_ and *s*_2_ are taken as −*v*_*E*,0_ and −*v*_*E*,1_, i.e. the negative of the excitatory mean voltage for each hemisphere; this is expected to be a reasonable LFP proxy ([47], [48], but see also [49]).

For a signal pair (*s*_1_, *s*_2_), we define the *symmetric* and *antisymmetric* parts as 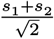 and 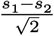, respectively. We define the *antisymmetric power fraction* of the signal pair as

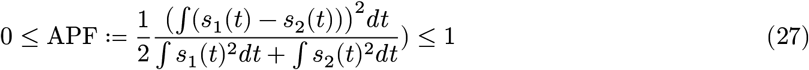

Note that this quantity evaluates to zero if *s*_1_ = *s*_2_, and is greater than 0 precisely when the solution to driven **4POP** dynamics is not constrained to lie in the symmetric subspace (the subspace ***x***_0_ = ***x***_1_, where ***x***_*j*_ denotes the concatenation of all state variables associated with hemisphere *j*).

Theta cycles are extracted from LFP by filtering signals in the 5-12 Hz band and identifying local extrema of the filtered signal; instantaneous phase estimates are obtained via the Hilbert transform. In contrast, the model output admits ground-truth theta cycles based on the phase of the sinusoidal input signal. Theta modulation index is defined as in [10], computed as the KL-divergence between the empirical (wavelet) amplitude distribution and a uniform distribution. For a circular sample *θ*_1_, …, *θ*_*N*_ *∈* [0, 2π), letting

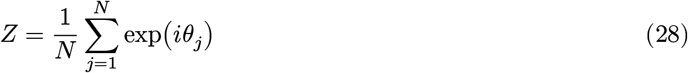

we define the circular mean as arg(*Z*) and the circular variance as 1 − |*Z*|.

To generate bifurcation diagrams, we have utilized a combination of HomotopyContinuation.jl ([50]; used to compute fold and Hopf loci) and MatCont ([51]; used to compute saddle node of limit cycles and Neimark-Sacker loci). First Lyapunov coefficients for Hopf points were computed manually ([52]). Maximal Lyapunov exponents were computed using the standard reorthonormalization procedure ([53]), as implemented in DynamicalSystems.jl ([54]). We have also utilized the stroboscopic map implementation in DynamicalSystems.jl to visualize chaotic attractors of the periodically forced dynamics. For numerical simulation of the microscopic spiking network dynamics, we utilize a standard Euler-stepping scheme, augmented by a reciprocal coordinate transformation (as described in [55]) to handle the voltage blowup at ∞. For numerical simulation of the macroscopic mean-field network dynamics, we utilize the Tsit5 scheme, as implemented in OrdinaryDiffEq.jl ([56]), through DynamicalSystems.jl. Finally, to perform the calculations described in the appendix, we utilized the computer algebra package Symbolics.jl ([57]).

### Experimental procedures

We reexamine recordings obtained and analyzed previously by Ghosh et al. ([10]). Briefly, the subset of data studied here was collected from head-fixed mice running on a ball. Linear probes were implanted to cross midline and obtain bilateral RSG field potentials. Full methodological details are found in [10], and all procedures were approved by the Institutional Animal Care and Use Committee at the University of Michigan.

## Results

### Oscillatory profile of RSG and model motivations

To illustrate the oscillatory phenomena of interest, we start with a brief reexamination of RSG extracellular recordings previously studied by Ghosh et al ([10]). Angled probes (Figure 1A) enabled simultaneous recording of field potentials from both hemispheres of mouse RSG during head-fixed running and sleep bouts. During active epochs and REM sleep, RSG LFP exhibits prominent background theta (4-12 Hz) oscillations, with superimposed high-frequency rhythms in the gamma (30-80 Hz) and spline (100-160 Hz) bands. Gamma and spline band power is modulated by theta phase, with the gamma/spline amplitude maximized during the rising/falling phases of theta, respectively (Figure 1C). Figure 1D depicts cross-spectral statistics of a typical channel pair extracted from superficial RSG channels across hemispheres; theta and gamma are in-phase, while splines are anti-phase.

**Figure 1:**
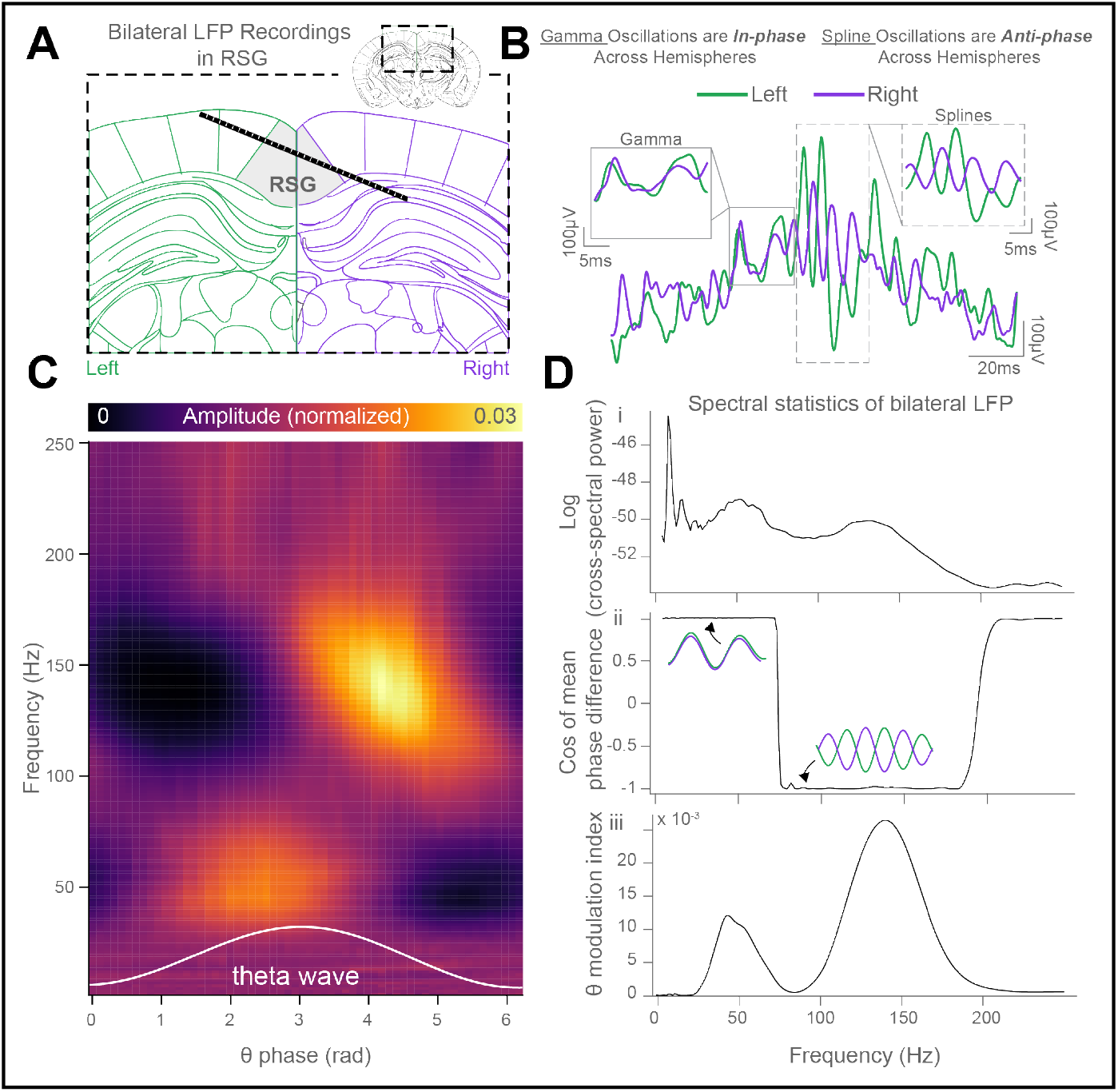
Theta-timescale alternation of in-phase gamma and anti-phase splines across retrosplenial hemispheres. ***A***, schematic of recording configuration. ***B***, schematic overview of experimental findings from Ghosh et al [10]; theta cycles recorded simultaneously from left and right RSG exhibit in-phase gamma on the rising phase and anti-phase splines on the falling phase. ***C***, phase-amplitude map averaged across theta cycles from a single recording. ***D***, spectral statistics from the same recording as in ***C***; i, logarithm of wavelet-cross spectral power, ii, cosine of the mean relative interhemispheric wavelet phase, iii, theta modulation index of high frequency oscillations.

Typical mechanisms for generating high-frequency oscillations in an excitatory-inhibitory circuit are the PING (pyramidal-interneuron gamma; [23]) and ING (interneuron gamma; [24]) instabilities, wherein oscillations are supported by excitatory → inhibitory → excitatory and inhibitory → inhibitory feedback loops, respectively. Because the PING loop is comprised of two synapses, while the ING loop requires only one, the latter mechanism generally supports higher-frequency oscillations than the former when the two loops coexist ([58], [38]). Indeed, both mechanisms can coexist in the same circuit, and experiments ([22]) suggest PING/ING mechanisms are likely responsible for production of low/high gamma oscillations, respectively, in dorsal subiculum. Although previous theoretical work has also suggested more elaborate mechanisms by which low and high gamma band oscillations could coexist in the same local circuit ([59], [60]), we elect to focus here on a relatively simple hypothesis: that RSG gamma and spline oscillations are locally (within a single hemisphere) accounted for by PING/ING mechanisms, respectively.

Subject to this assumption, our goals are therefore 1) to construct model interhemispheric networks for which inphase PING (IP-PING) and anti-phase ING (AP-ING) oscillations can coexist, 2) to characterize the possible dynamics of these networks, with and without extrinsic theta rhythmic forcing, and 3) to understand mathematically which model features control the simultaneous cross-hemispheric phase relationship for PING and ING oscillations.

To accomplish these goals, we perform numerical experiments within the established modeling framework of exact neural mass models derived from networks of globally-coupled quadratic integrate-and-fire (QIF) neurons ([28], [25], [30], [61]). We consider networks comprised of interacting homogeneous populations of QIF neurons, each receiving a Cauchy white noise input ([35], [30]); remarkably, dynamics of such networks are exactly captured by a low-dimensional set of ordinary differential equations (**QIF-NM**, Equation 9). Importantly, the process of deriving mean-field equations from the microscopic dynamics averages over the neurons in each population (Equation 2 and Equation 3), and not over time; thus, the **QIF-NM** model retains fine timing details of population dynamics, while discarding the timing of individual constituent spikes. Together, low-dimensionality and exactness of the mean-field dynamics enable use of numerical bifurcation theory ([52]) to study key dynamical behaviors of the network, such as population oscillations, in both strongly and weakly synchronized regimes. We supplement these numerical studies of **QIF-NM** networks with an analytical study of simplified firing rate models (Analytical Appendix), providing evidence that the results and underlying mechanisms are qualitatively robust to the choice of modeling framework, and a deeper understanding of precisely which model parameters control the oscillatory regimes of interest.

To illustrate the **QIF-NM** framework, we first consider a model network capable of generating both PING and ING oscillations: this is the **2POP** motif, comprised of one excitatory (E) and one inhibitory (I) population (Equation 23). Based on the results of previous patch-clamp studies ([41],[7], [42]), we assume 1) there is little if any recurrent excitation in superficial RSG and 2) the decay timescale of excitatory inputs onto inhibitory neurons is shorter than the decay timescale of inhibitory inputs onto excitatory neurons. With the network regime further constrained by these assumptions, Figure 2 depicts a typical dynamical landscape for the **2POP** network, upon variation of distinguished parameters *η*_*E*_ and *η*_*I*_ . These parameters describe the level of external DC (constant in time) drive to the E and I populations, respectively. When (*η*_*E*_, *η*_*I*_ ) is chosen to lie in the shaded grey region (Figure 2B), the network settles into a stable asynchronous steady state (Figure 2D, middle). Accordingly, we call the shaded grey area the *stability region*. The boundary of the stability region is generally comprised by a union of *local bifurcation curves*, each of which corresponds to a particular mathematical mechanism by which stability of the asynchronous state is lost. In the case of **2POP**, the boundary of the stability region is a single local bifurcation curve, called a *Hopf curve*.

**Figure 2:**
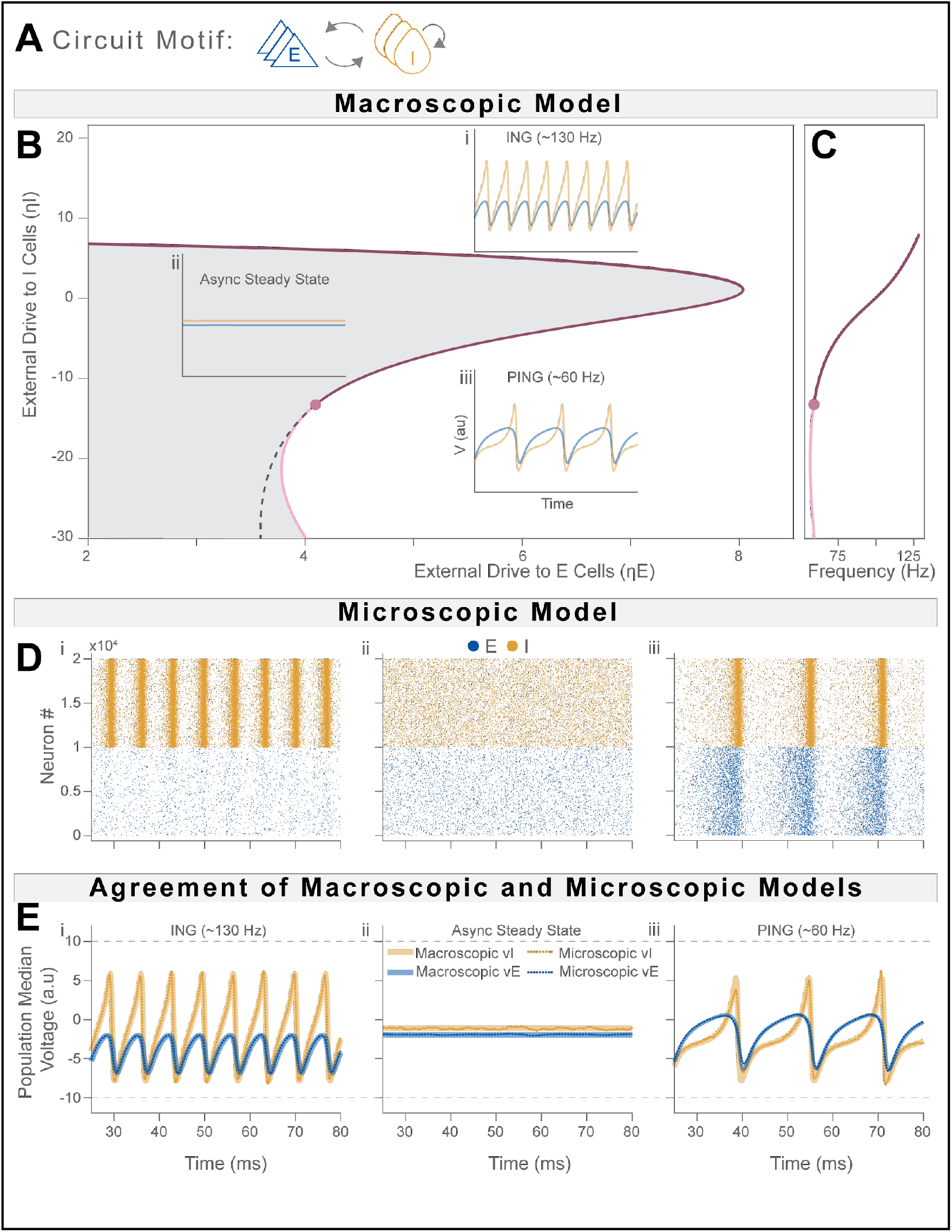
Dynamics of two population (**2POP**) excitatory-inhibitory network. ***A***, diagram of **2POP** circuit motif; coupled E and I populations, with no recurrent excitation among E cells. ***B***, two-parameter bifurcation diagram in (*η*_*E*_, *η*_*I*_ ) external drive plane; dark red: supercritical Hopf, light red: subcritical Hopf, dashed black: saddle-node of limit cycles (bounding region of asynch/PING bistability). ***C***, frequency of bifurcating eigenvalue along Hopf curve, tracking the transition from PING to ING-dominated regime upon varying *η*_*I*_. ***D***, raster plots of microscopic network simulation results, depicting ING/asynch/PING dynamics. ***E***, median voltage across cells from microscopic simulations shown in, with mean-field predictions overlaid; note the close agreement. Results displayed utilize *η*_*E*_ = 6.5 and *η*_*I*_ = 10 (ING), *η*_*I*_ = 0 (asynchronous), *η*_*I*_ = −10 (PING). Microscopic networks consist of *N*_*E*_ = *N*_*I*_ = 10^4^ neurons in excitatory and inhibitory populations, respectively. All other parameters are taken from the default set (see Methods).

Varying (*η*_*E*_, *η*_*I*_ ) to cross the Hopf curve gives rise to synchronous macroscopic oscillations. For a particular range of *η*_*E*_ values (around *η*_*E*_ *∈* (4, 8) in Figure 2B), if we fix *η*_*E*_ and vary only *η*_*I*_, the asynchronous state can lose stability in two qualitatively distinct ways. At high *η*_*I*_, the network produces ING-like oscillations characterized by higher frequency and strong hyperpolarization of the excitatory population; these oscillations may persist upon ablation of feedforward excitation (setting the E → I synaptic weight to zero). In contrast, at low *η*_*I*_, the network produces PING-like oscillations, characterized by lower frequency and relative depolarization of the excitatory population; these PING-like oscillations are instead extinguished upon the ablation of feedforward excitation. Despite the qualitative distinction, we emphasize that the transition between ING and PING is continuous along the Hopf boundary (Figure 2C), and these solutions exist on the same branch of periodic orbits. This bifurcation structure anticipates that a slow variation of *η*_*I*_, back and forth across the stability region, can result in transit between ING and PING rhythms, with the difference in their frequencies modulated by the fixed background value of *η*_*E*_. Note that Figure 2 depicts the close agreement between the microscopic network simulations (Figure 2D) and the predictions of the macroscopic network equations (Figure 2E); this justifies our focus, in what follows, to exclusively analyzing dynamics of the macroscopic model.

Before proceeding, we note that although the results presented in Figure 2 directly reflect the dynamical landscape of **QIF-NM 2POP** circuits, simplified firing rate networks exhibit an analogous oscillatory landscape (Analytical Appendix, Figure S1, Figure S3), indicating qualitative features of the architectural motifs examined here are indeed expected to generalize across modeling frameworks (previously demonstrated also in [38], by similar analyses).

### Dynamical landscape of symmetrically coupled E-I modules

We model each hemisphere as a **2POP** pair capable of locally generating PING and ING oscillations; IP-PING and AP-ING may then occur when two **2POP** pairs are coupled into a **4POP** motif via delayed long-range projections. The **4POP-IE** model (Figure 3A) takes all interhemispheric projections to be from ipsilateral E-cells to contralateral I-cells. This motif is representative of the situation where interactions between I populations across hemispheres are mediated exclusively via the E population which drives local PING oscillations. In contrast, the **4POP-II** model (Figure 4A) takes the only nonzero interhemispheric projections to be from ipsilateral I-cells to contralateral I-cells. This scenario represents the case where I-cell populations interact via some pathway which is independent of the E-cell population driving local PING. Projections of the ipsi-E to contra-I type are common, and known to exist for both granular ([7], [42]) and dysgranular ([62]) retrosplenial cortices. Projections of the ipsi-I to contra-I type are rarer, but known to exist ([63], [64]) and frequently neglected in modeling studies.

**Figure 3:**
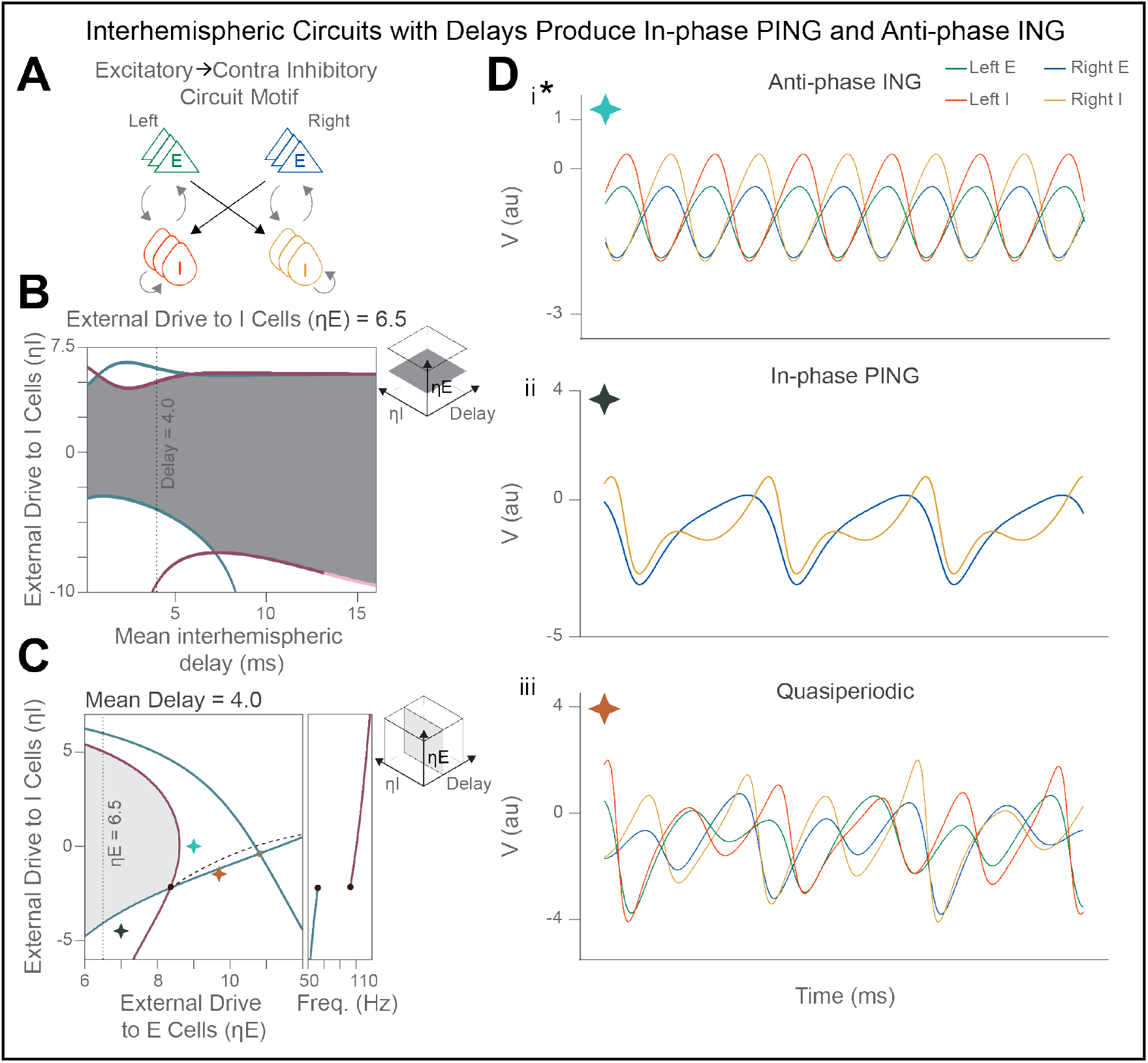
Dynamics and two-parameter bifurcation diagrams for the **4POP-IE** model with *ρ*_*IE*_ = 0.5. ***A***, circuit diagram of the **4POP-IE** motif, comprised of two **2POP** pairs coupled with cross-hemispheric **E → I** projections. ***B***, bifurcations induced by varying drive to inhibitory population (*η*_*I*_ ) and cross-hemispheric delay. Dark blue: supercritical longitudinal Hopf, light blue: subcritical longitudinal Hopf, dark red: supercritical transverse Hopf, light red: subcritical transverse Hopf; around *D* = 4 ms, IP-PING and AP-ING coexist. ***C***, bifurcations in external drive to excitatory and inhibitory populations, with fixed mean cross-hemispheric delay (*D* = 4 ms). Same color scheme as in, with dashed Neimark-Sacker curve emerging from double Hopf point; right depicts frequency of bifurcating eigenvalue along boundary of stability region (grey). ***D***, example dynamics of **4POP-IE**, corresponding with drive parameters chosen according to star markers in ***C***. Note in ii, the apparent presence of only two traces (right E and I) reflects the precise symmetry between the two hemispheres; all four traces are present and perfectly in-phase.

**Figure 4:**
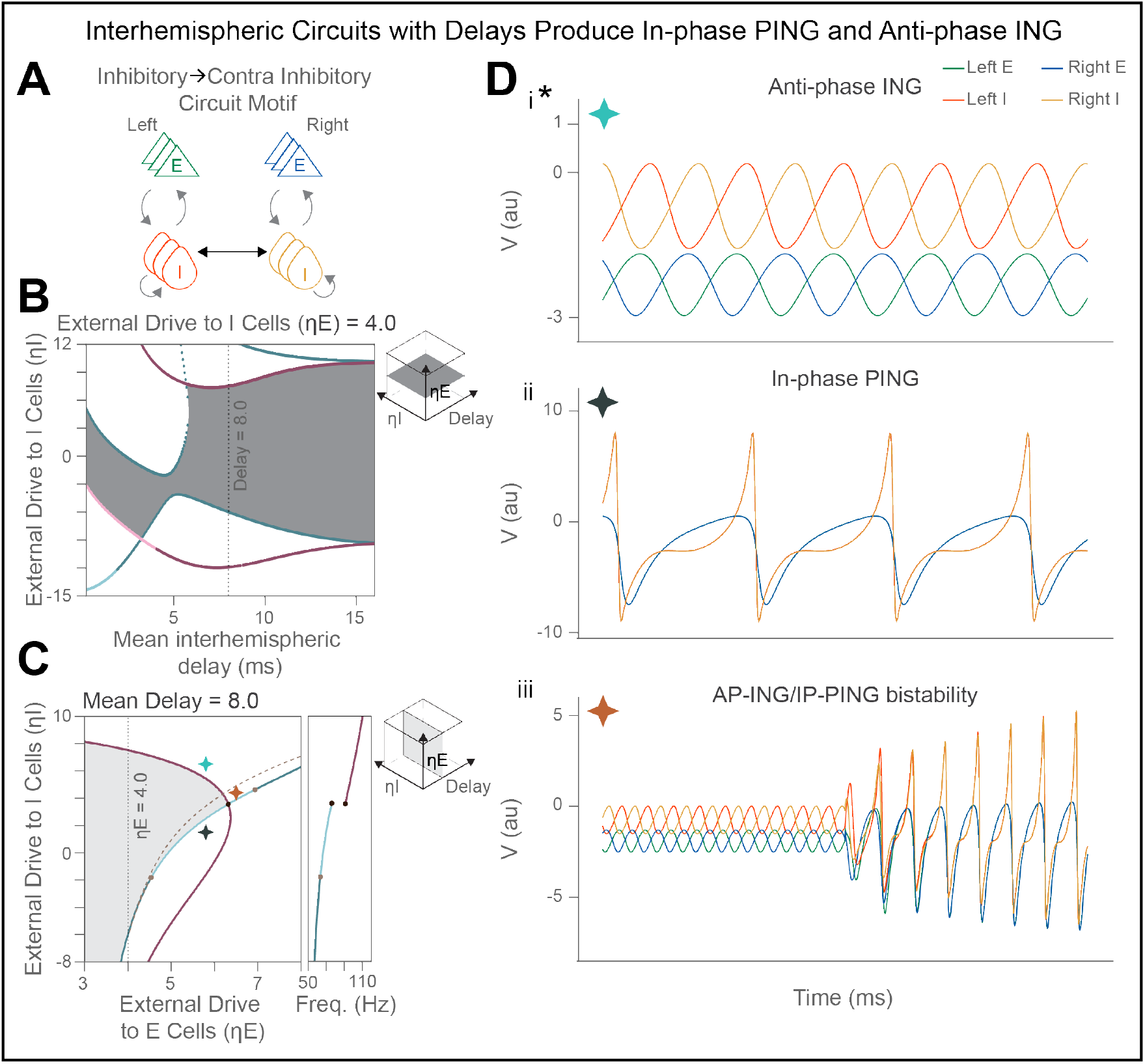
Dynamics and two-parameter bifurcation diagrams for the **4POP-II** model with *ρ*_*II*_= 0.5. ***A***, circuit diagram of the **4POP-II** motif, comprised of two **2POP** pairs coupled with cross-hemispheric **I → I** projections. ***B***, bifurcations induced by varying drive to inhibitory population (*η*_*I*_ ) and cross-hemispheric delay. Color scheme for bifurcations identical to that in Figure 3; around *D* = 8 ms, IP-PING and AP-ING coexist., bifurcations in external drive to excitatory and inhibitory populations, with fixed mean cross-hemispheric delay (*D* = 8 ms). Same color scheme as in ***B***, with with dashed saddle-node of limit cycles emerging from generalized Hopf point; right depicts frequency of bifurcating eigenvalue along boundary of stability region (grey). ***D***, example dynamics of **4POP-II**, corresponding with drive parameters chosen according to star markers in ***C***. As in Figure 3, the inphase panel (ii) depicts perfect synchronization across hemispheres.

Both **4POP** models support a symmetric steady state, wherein the firing rate of each population is exactly equal to that of its interhemispheric counterpart. As in the **2POP** model, this symmetric steady state can lose stability upon variation of external DC drives. In symmetric dynamical systems, such as the **4POP** model, it is possible to classify these instabilities of the symmetric steady state into precisely two types: *longitudinal* and *transverse*. Longitudinal instabilities are essentially those which are already present in the **2POP** model, while transverse instabilities exist only in the full **4POP** model and give rise to network states that are symmetry-broken in space or time. In particular, longitudinal Hopf bifurcations give rise to collective oscillations which are *in-phase* across hemispheres, while transverse Hopf bifurcations give rise to oscillations which are *anti-phase* across hemispheres. Therefore, in analogy with previous observations from the **2POP** model, a sufficient condition for coexistence of IP-PING and AP-ING is the presence of a longitudinal Hopf bifurcation at low values of *η*_*I*_, and a transverse Hopf bifurcation at higher values of *η*_*I*_ . We emphasize this is distinct from the notion of bistability, wherein both oscillations are attainable for a single fixed parameter set, via distinct initial conditions (although we have also observed bistability between IP-PING and AP-ING; Figure 4D/iii).

With our default parameter set chosen (see Methods), we fix *η*_*E*_ and examine bifurcations of the symmetric equilibrium in the (*D, η*_*I*_ ) parameter plane (Figure 3B, Figure 4B), where *D* denotes the mean interhemispheric delay. Both the **4POP-IE** and **4POP-II** models admit a range of values *D* for which this equilibrium loses stability at high *η*_*I*_ to a transverse Hopf (around *D* ≈ 2-5 ms, yielding AP-ING) and at low *η*_*I*_ to a longitudinal Hopf (around *D* ≈ 6-10 ms, yielding IP-PING). Exactly these parameter ranges furnish the model regimes most consistent with experimentally observed RSG spectral content. Accordingly, to explore the model dynamics in more detail within these regimes, we fix the mean interhemispheric delay for **4POP-IE** at *D* = 4 ms and for **4POP-II** at *D* = 8 ms.

An additional important parameter for each **4POP-XY** model is its *interhemispheric weight ratio, ρ*_*XY*_ *∈* [0, 1]. This represents the fraction of total projection strength which is directed towards the contralateral, rather than ipsilateral, hemisphere. For example, *ρ*_*IE*_ = 0 describes the **4POP-IE** model in which all E → I projection weight is local (i.e. the degenerate case of two uncoupled **2POP** pairs), while *ρ*_*IE*_ = 1 describes the model for which all E → I projection weight is long-range, with no local feedforward inhibition.

Figure 3C and Figure 4C depict the (*η*_*E*_, *η*_*I*_ ) plane for the models with *ρ*_*IE*_ = *ρ*_*II*_ = 0.5, respectively, where local and long-range projections are of equal strength. In both **4POP** scenarios, the single Hopf curve bounding the stability region for **2POP** is perturbed into distinct transverse and longitudinal curves, which meet at what is called a *double Hopf point*. It is important to note that Hopf bifurcations may occur in two forms: *supercritical* and *subcritical*. At supercritical Hopf bifurcations, the transition from steady state to oscillatory dynamics occurs smoothly, with the oscillation amplitude shrinking cleanly to zero as one approaches the Hopf curve from the oscillatory side. At subcritical Hopf bifurcations, the observed oscillation amplitude may jump abruptly at the Hopf curve. Note that in **4POP-IE**, all Hopf points bounding the stability region are supercritical, while in **4POP-II**, the symmetric equilibrium can undergo a subcritical longitudinal Hopf bifurcation.

Just as steady states can undergo bifurcations giving rise to new dynamics (such as collective oscillations), oscillatory solutions can undergo bifurcations giving rise to more complex *quasiperiodic* solutions, which exhibit activity reflecting multiple distinct frequencies. This occurs along what is called a *Neimark-Sacker* curve . In **4POP-IE**, a Neimark-Sacker curve emerges from the double Hopf point, beyond which the AP-ING limit cycle bifurcates to a quasiperiodic attractor with mixed ING-PING dynamics.

In **4POP-II**, the *generalized Hopf point* (where the longitudinal Hopf bifurcation switches from supercritical to subcritical) on the stability boundary emits a *saddle-node of limit cycles*, a curve along which stable and unstable periodic solutions may collide and annihilate one another. This curve predicts the existence of a regime of bistability between AP-ING and IP-PING, which we confirm with numerical simulations. Indeed, in this parameter regime, numerical experiments indicate that varying *η*_*I*_ modulates the size of the basin of attraction for each oscillation type (data not shown).

We note that similarly to the isolated **2POP** circuit, threshold-linear rate networks with **4POP-IE** and **4POP-II** architectures exhibit a Hopf landscape which is qualitatively analogous to the one described here for **QIF-NM 4POP** networks (Analytical Appendix, Figure S4; but note that the criticality of the Hopf bifurcations depends on the nonlinear dynamical features, and cannot be expected to generalize across models).

Next, we examine the effect of varying *ρ*_*XY*_ from 0 to 1 (Figure 5), crucial to understand how the strength of cross-hemispheric projections affects network dynamics. In the **4POP-IE** model (Figure 5B), for all sufficiently small *ρ*_*IE*_, we observe the same scenario as described above for *ρ*_*IE*_ = 0.5; namely, the stability boundary is comprised of supercritical transverse and longitudinal Hopf points. Upon increasing *ρ*_*IE*_ further, an additional curve of transverse *fold bifurcations* comes in from *η*_*I*_ = −∞, such that it is positioned close to the IP-PING longitudinal Hopf curve. This transverse fold (also called a *pitchfork* in the present symmetric setting) curve indicates a third distinct instability of the symmetric equilibrium, for sufficiently large *ρ*_*IE*_: instead of giving rise to an oscillation, the symmetric equilibrium gives rise to two stable, symmetry-broken steady states, for which the two hemispheres have distinct steady state voltages (Figure 5B/iv, teal marker). The **4POP-II** model (Figure 5C) also exhibits the symmetry-breaking transverse fold bifurcation for *ρ*_*II*_ large; however, in this case, the curve comes in from *η*_*I*_ = +∞ (Figure 5C/iii). Yet another difference is that as *ρ*_*II*_ increases, the longitudinal Hopf of **4POP-II** changes criticality around *ρ*_*II*_ = 0.5, and then grows to encompass the entire stability region (Figure 5C/iv).

**Figure 5:**
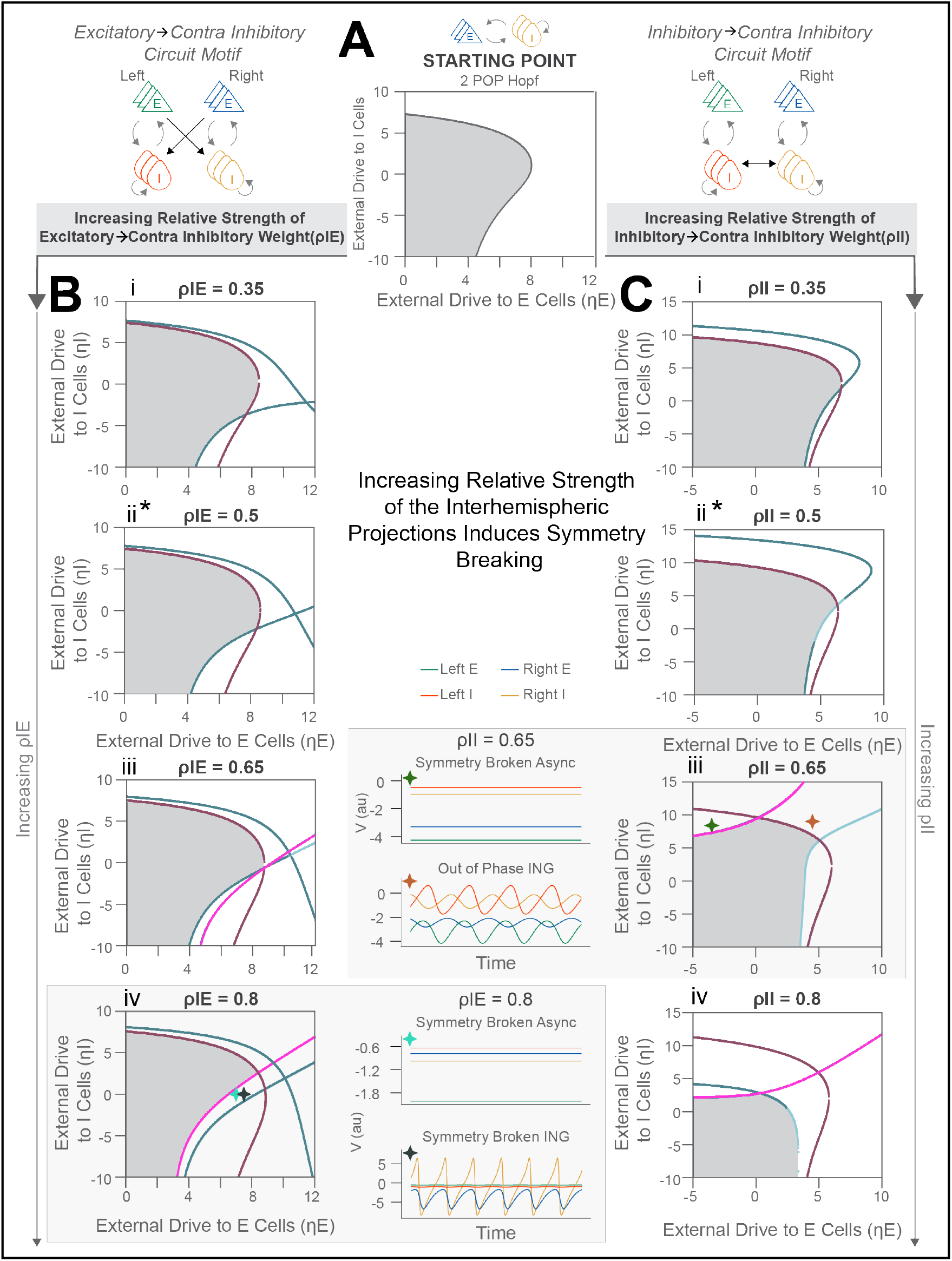
Impact of varying the interhemispheric weight ratio in **4POP-IE/II. *A***, Two-parameter bifurcation diagram in (*η*_*E*_, *η*_*I*_ )-plane, depicting Hopf curve for **2POP** model as in Figure 2B. ***B***, two-parameter bifurcation diagrams (in external drive plane) for **4POP-IE**, upon variation of *ρ*_*IE*_; same color scheme as in Figure 3. At large *ρ*_*IE*_, the symmetric steady state undergoes a transverse fold bifurcation (pink curves), giving rise to two symmetry-broken steady states; the symmetry broken steady state can itself undergo Hopf bifurcation, yielding symmetry-broken ING (shown) and symmetry-broken PING (not shown) oscillations. ***C***, same as ***B*** for **4POP-II** motif. In **4POP-II** with *ρ*_*II*_ = 0.65, in addition to emergence of symmetry-broken equilibria, we observe a bistable pair of limit cycles exhibiting out-of-phase ING, suggestive that the anti-phase ING limit cycle itself directly undergoes pitchfork bifurcation at high *η*_*I*_ . We note the distinct locations of the transverse fold curves across the **4POP-IE** and **4POP-II** motifs, which gives rise to phase-locked symmetry breaking under periodic forcing (Figure S7).

To conclude our description of the **4POP-XY** dynamical landscapes, we note the possibility of further symmetry-broken dynamics that can exist at high *ρ*_*XY*_ . In both **4POP-XY** models, the symmetry-broken equilibrium at high *ρ*_*XY*_ can itself undergo Hopf bifurcation (Figure 5B/iv, black marker; Figure 5C/iii, gold marker), giving rise to symmetry-broken limit cycles, wherein one hemisphere oscillates and the other is nearly asynchronous. For large *ρ*_*II*_ ≈ 0.65, increasing *η*_*I*_ further beyond the transverse Hopf curve can transition the anti-phase oscillations into out-of-phase oscillations, where the phase difference of ING rhythms across hemispheres is close to, but distinct from, π (Figure 5C/iii, gold marker); numerical evaluation of the *Floquet exponents* (a way to quantify stability of periodic solutions) suggests these out-of-phase ING oscillations emerge via a *pitchfork bifurcation of limit cycles* (data not shown). The phase deviation from π increases monotonically with *η*_*I*_, and thus vicinity to the pitchfork of limit cycles provides a mechanism that can be exploited by biological neural networks to precisely modulate the relative interhemispheric phase.

### Switching between in-phase PING and anti-phase ING via external theta inputs

Having identified two model networks, **4POP-IE** and **4POP-II**, for which AP-ING and IP-PING coexist, we next examine the network dynamics under theta-rhythmic AC forcing. To mimic GABAergic septal inputs onto RSG I cells ([8]), we apply a harmonic forcing with period 150 ms to the inhibitory populations of both hemispheres. These theta-rhythmic inputs are assumed synchronous across hemispheres, in line with experimental observations ([12], [10]). Accordingly, throughout what follows we refer to the phase of the theta rhythm at which inhibitory neurons are most silenced as the *trough* of theta, and the phase at which they are most active as the *peak*. We call the trough-to-peak interval of each cycle its *rising phase*, and the peak-to-trough interval its *falling phase*. In this section, we use the default parameters with forcing amplitude *A*_*I*_ = −9; however, we continue to vary the DC drive parameters (*η*_*E*_, *η*_*I*_ ) and interhemispheric weight ratios *ρ*_*XY*_ .

Since we utilize theta forcing which is precisely in-phase across hemispheres, sufficiently low amplitude modulation of the symmetric equilibrium results in a *symmetric limit cycle solution*; and similarly to the equilibrium case, this symmetric limit cycle may itself undergo transverse or longitudinal bifurcations. To characterize where in the (*η*_*E*_, *η*_*I*_ ) plane these symmetric limit cycles lose stability, we vary the DC drive parameters and compute the antisymmetric power fraction from an LFP proxy signal (APF; see Methods); this solution measure vanishes only when both hemispheres are precisely in-phase, thus localizing bifurcations which redistribute solution power into the antisymmetric subspace. Indeed, in both **4POP-XY** models, we observe a sharp transition from zero to nonzero APF in the (*η*_*E*_, *η*_*I*_ ) plane, whose contour is traced by a transverse Neimark-Sacker curve (Figure 6B, Figure 7B). Beyond this curve, the symmetric limit cycle bifurcates into quasiperiodic trajectories which exhibit phase-amplitude coupled AP-ING bursts (Figure 6D, Figure 7D, i). In **4POP-II**, we also find a longitudinal Neimark-Sacker curve (Figure 7B/C), yielding phase-amplitude coupled IP-PING bursts (Figure 7D, iii). Hunting next for solutions which exhibit PING/ING alternation, we find that **4POP-IE** supports symmetric limit cycle solutions which exhibit nested IP-PING/IP-ING alternation (Figure 6D, ii); then, at higher values of *η*_*I*_, just beyond the transverse Neimark-Sacker curve, we note quasiperiodic solutions with IP-PING/AP-ING alternation. The **4POP-II** model also supports IP-PING/AP-ING alternation (Figure 7D, ii), but these switching solutions are instead *chaotic*; they exhibit nested IP-PING/AP-ING oscillations whose amplitudes vary irregularly across theta cycles. We quantify the extent of this chaos in Figure 7C using the *maximum Lyapunov exponent* ([65]), which measures the rate at which nearby dynamical trajectories diverge from one another. These dynamics presumably reflect transit via theta inputs through the bistable regime observed in Figure 4D, iii. Note that across both **4POP-XY** models, IP-PING and AP-ING consistently occur during the rising and falling phases, respectively, of the driving theta rhythm, reflecting the geometric configuration of the bifurcation curves in the **2POP** parameter plane.

**Figure 6:**
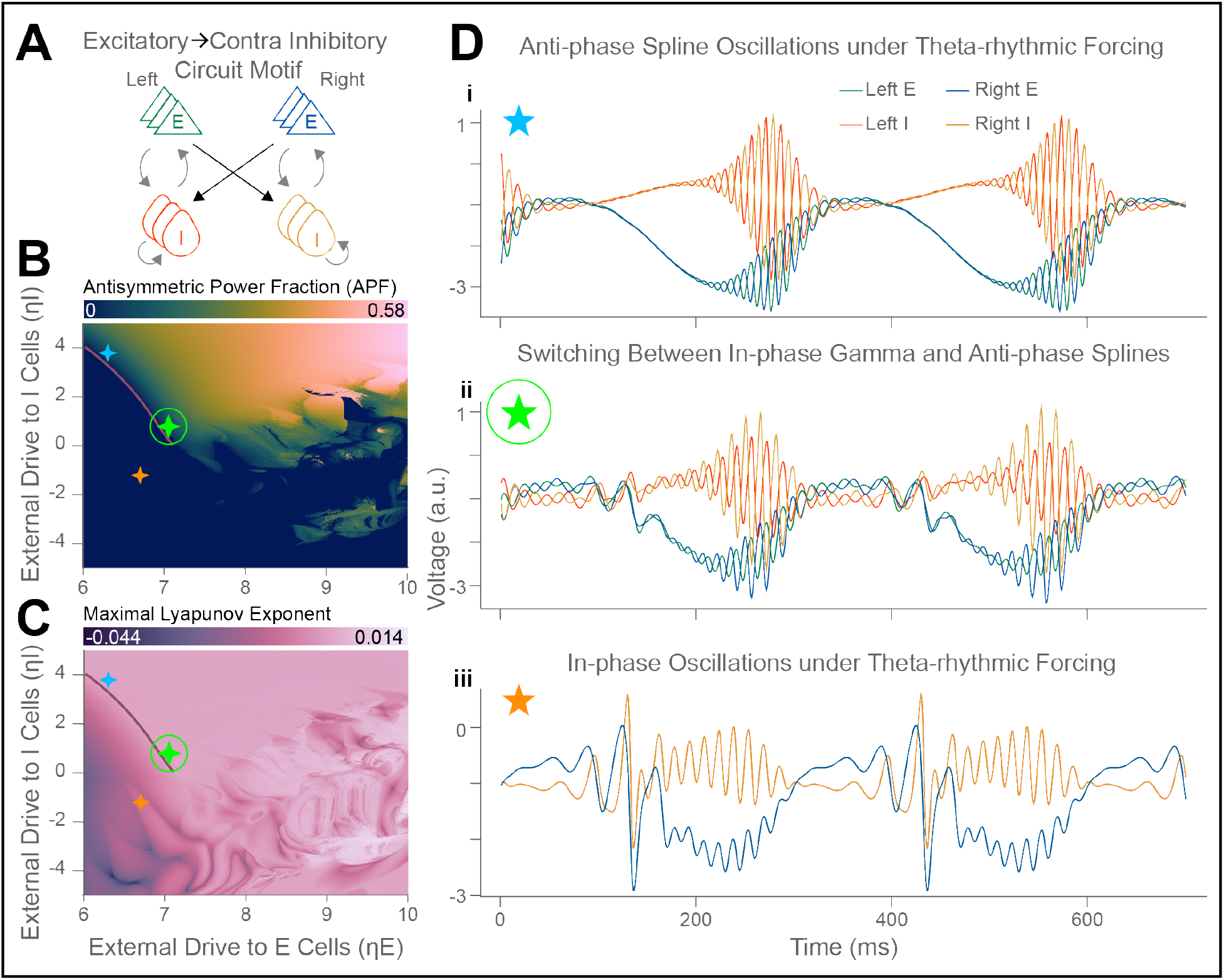
Dynamics of theta-forced **4POP-IE** network. ***A***, circuit diagram of the **4POP-IE** motif. ***B***, antisymmetric power fraction (APF) of forced **4POP-IE** solution as (*η*_*E*_, *η*_*I*_ ) varies, indicating where symmetric trajectories lose stability. ***C***, maximum Lyapunov exponent of forced **4POP-IE** solution. Red curve in ***B*** and ***C*** depicts transverse Neimark-Sacker bifurcation of the symmetric theta-rhythmic limit cycle solution. Note that the maximum Lyapunov exponent across the entire considered (*η*_*E*_, *η*_*I*_ ) parameter plane is bounded above by 0.014; hence, when the forced **4POP-IE** is chaotic, it is weakly so. ***D*, i**. Quasiperiodic trajectory emerging from the transverse Neimark-Sacker curve; this solution typifies phase-amplitude coupled AP-ING oscillations. **ii**. At lower *η*_*I*_, low amplitude IP-PING oscillations alternate with higher amplitude AP-ING oscillations, producing RSG-like patterning (circled green star). **iii**. Left of the transverse Neimark-Sacker curve, symmetric limit cycle solutions can develop apparent in-phase PING-ING alternation, reflecting the ability of **2POP** itself to achieve this behavior. Parameters used to generate this figure are *A*_*I*_ = 9, *ρ*_*IE*_ = 0.6, *D* = 4 ms.

**Figure 7:**
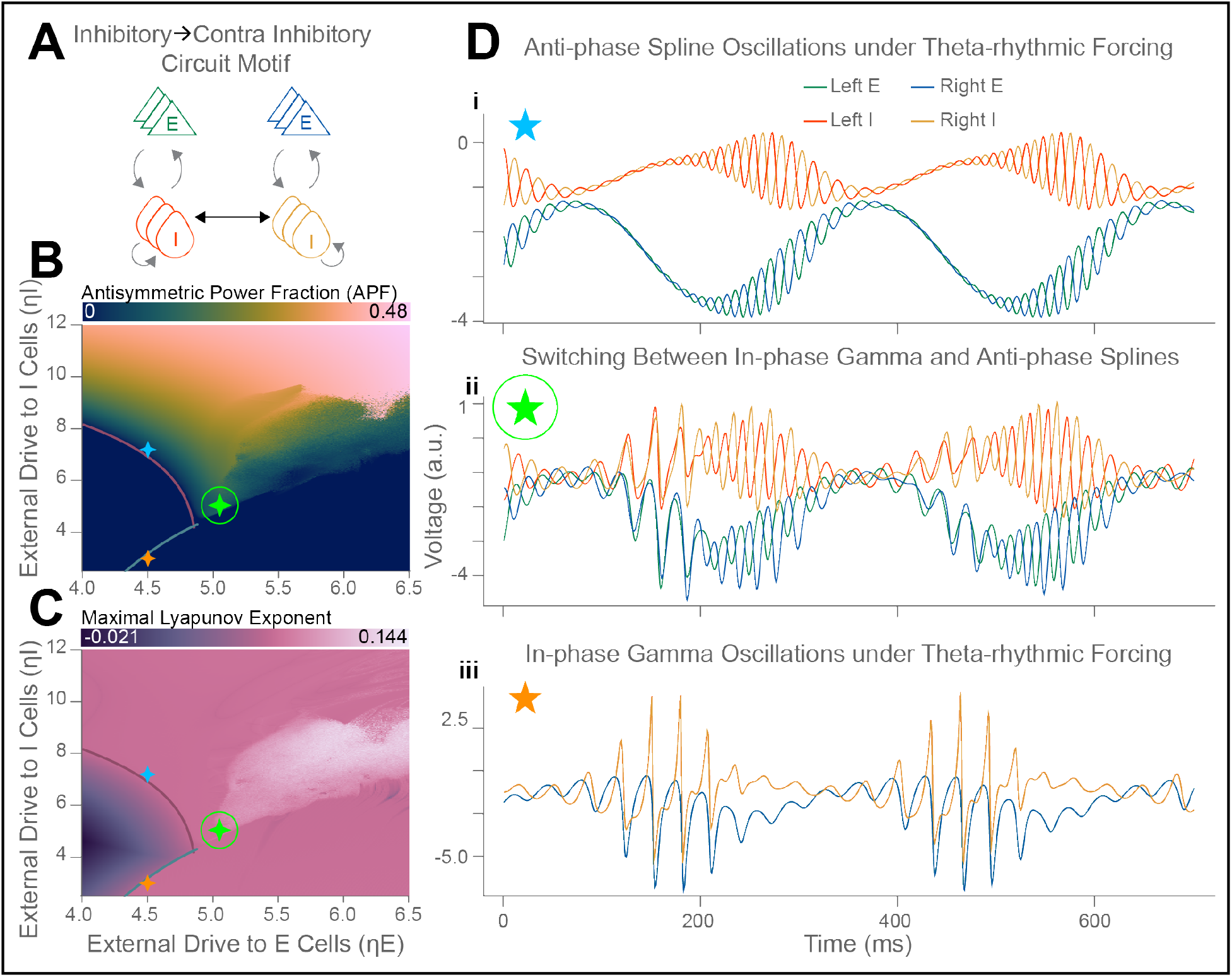
Dynamics of theta-forced **4POP-II** network. ***A***, circuit diagram of the **4POP-II** motif. ***B***, antisymmetric power fraction (APF) of forced **4POP-II**, as in Figure 6. ***C***, maximum Lyapunov exponent of forced **4POP-II** solution. Note the broad region of chaos at higher *η*_*E*_, with comparatively high chaoticity relative to **4POP-IE**. Red curve in ***B*** and ***C***: transverse Neimark-Sacker bifurcation of the symmetric theta-rhythmic limit cycle solution. Blue curve in ***B*** and ***C***: longitudinal Neimark Sacker bifurcation. ***D***, **i**. As in **4POP-IE**, the transverse Neimark-Sacker bifurcation of the symmetric theta-rhythmic limit cycle gives rise to quasiperiodic solutions exhibiting phase-amplitude coupled AP-ING oscillations. **ii**. Beyond the double Neimark-Sacker point where the two curves intersect, we observe chaotic trajectories exhibiting RSG-like AP-ING/IP-PING alternation. **iii**. Unlike in **4POP-IE, 4POP-II** exhibits also a longitudinal Neimark-Sacker bifurcation (blue curve), yielding phase-amplitude coupled IP-PING oscillations. Parameters used to generate this figure are *A*_*I*_ = 9, *ρ*_*II*_ = 0.65, *D* = 8 ms.

To examine the robustness of these dynamics in the (biologically unavoidable) presence of noise, we considered also a “mesoscopic” neural mass model, which approximates the dynamics of finite size QIF networks by incorporating a multiplicative noise term into the macroscopic mean field equations (see Methods). Indeed, Figure S7A,Bi depicts mesoscopic solutions to both **4POP-XY** networks which express noisy IP-PING/AP-ING alternation, demonstrating both network motifs produce RSG-like dynamics when subjected to stochastic drive. At larger *ρ*_*IE*_, the mesoscopic model exhibits *transient symmetry-breaking* (TSB) events (Figure S7Aii-iii), predicted by emergence of the transverse fold curve in Figure 5B. These TSB events appear as deviations of the mean voltage across hemispheres (Figure S7Aii-iii), and contribute theta-band power to the antisymmetric part of the signal pair. Consistent with the positioning of this transverse fold curve, TSB events in **4POP-IE** tend to occur during the rising phase of theta, i.e. shortly after *η*_*I*_ is lowest, and simultaneously with IP-PING bursts. In contrast, TSB events in **4POP-II** occur at high values of *η*_*I*_, during the falling phase of the theta input and simultaneously with AP-ING (Figure S7Bii-iii). At high *ρ*_*IE*_, TSB events can evoke symmetry-broken ING bursts (Figure S7Aiii, as in Figure 5B), in which oscillation amplitude markedly differs across hemispheres. At high *ρ*_*II*_ and low *η*_*E*_, we also observe out-of-phase ING oscillations (Figure S7Biii, as in Figure 5C). Thus, both models support theta-timescale competitive dynamics, which may exist alongside phase-amplitude coupled IP-PING/AP-ING oscillations; it would be interesting to hunt for putative signatures of this dynamic in experimental recordings.

Lastly, we directly compare spectral statistics generated by the mesoscopic neural mass model with those obtained from experimental data. Wavelet cross-spectra of both **4POP-XY** models exhibit two distinct spectral peaks at high frequencies, corresponding with PING and ING (Figure 8G/I). Circular statistics of the relative interhemispheric phase indicate that PING and ING across hemispheres are indeed in-phase and anti-phase, respectively (Figure 8G/I, insets).

**Figure 8:**
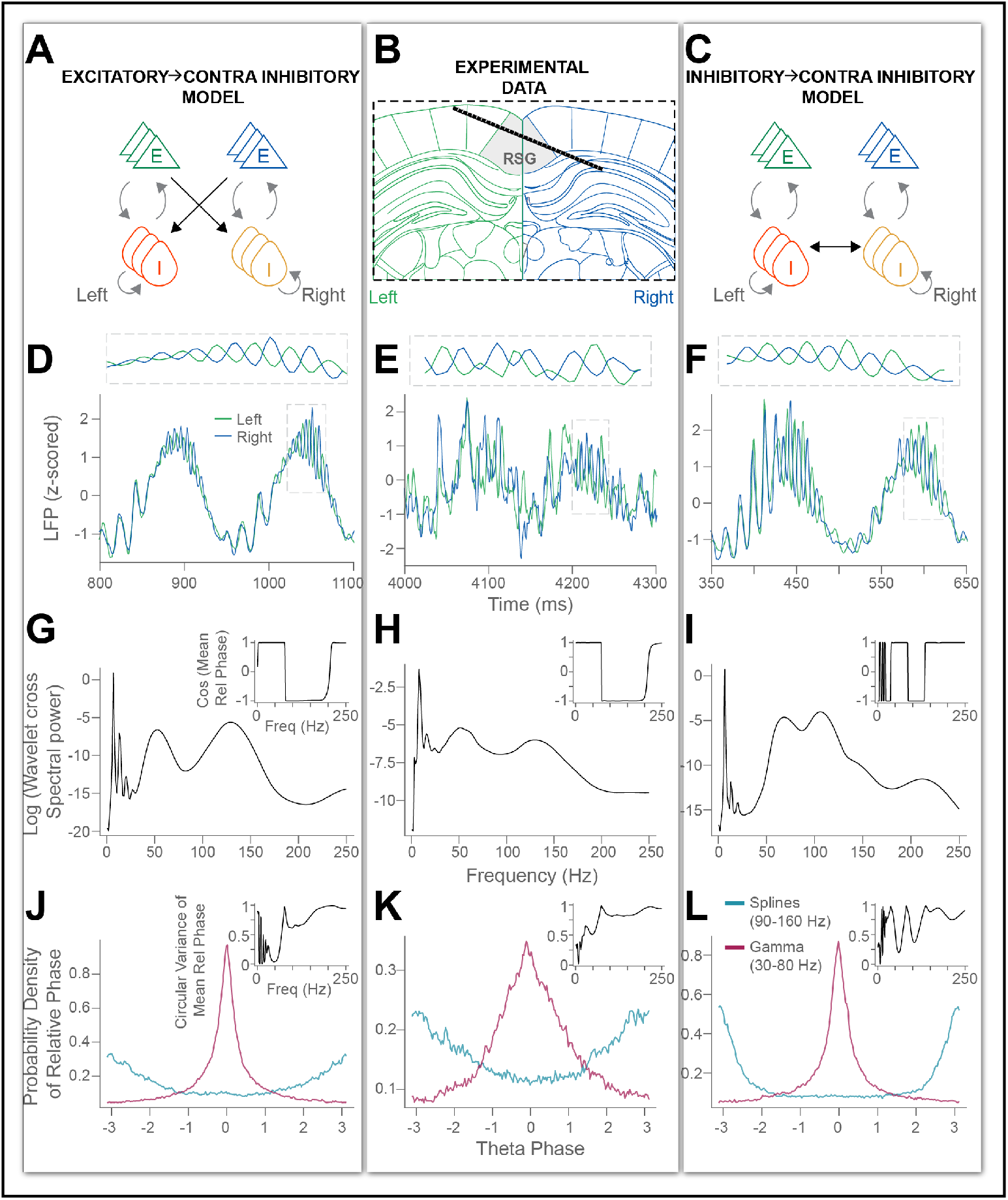
Comparison of spectral statistics across experiment and model. ***A***, circuit motif for **4POP-IE** model. ***B***, example probe trajectory for bilateral RSG recordings. C, circuit motif for **4POP-II** model. ***D***, two example theta cycles from mesoscopic **4POP-IE** model with (*η*_*E*_, *η*_*I*_) = (5, −0.8), *ρ*_*IE*_ = 0.6, *A*_*I*_ = 9, *N*_*E*_ = *N*_*I*_ = 8000. ***E***, two example theta cycles from experimental recording. F, two example theta cycles from mesoscopic **4POP-II** model with (*η*_*E*_, *η*_*I*_) = (3.5, 2.0), *ρ*_*II*_ = 0.65, *A*_*I*_ = 9, *N*_*E*_ = *N*_*I*_ = 8000. ***G-I***, wavelet cross-spectra averaged over 1600 seconds of synthetic data for **4POP** models, and 1000 seconds of experimental data (active exploration only); main plots depict logarithm of cross-spectral power, while insets show cosine of the mean interhemispheric relative phase. ***J-L***, histograms of the relative interhemispheric Hilbert phase extracted from broadband spline (90-160 Hz) and gamma (30-80 Hz) signals; insets quantify precision of interhemispheric phase-locking via circular variance of the relative phase.

We emphasize several features of the comparison between models and data. First, we observe that in the spline/ AP-ING band, experimental recordings indicate a fairly high circular variance of the relative interhemispheric phase (variance ≈ 0.8, Figure 8K, inset), and a broad range of frequencies in which this variance is reduced from the null value of 1. For the chosen parameter settings, this fairly variable cross-hemispheric phase difference and broadband nature of the spectral peaks are better recapitulated by the **4POP-IE** (variance ≈ 0.6, Figure 8J, inset) than by **4POP-II** (variance ≈ 0.5, Figure 8L, inset). Furthermore, we note that the spectral peaks are closer to one another in the **4POP-II** model (≈ 40 Hz, Figure 8I) than in the **4POP-IE** model (≈ 80 Hz, Figure 8G), with the **4POP-IE** more closely reproducing the frequency difference observed *in vivo* (≈ 90 Hz, Figure 8H). In fact, this discrepancy between the two motifs is reproduced and explained by direct mathematical analysis of simplified firing rate models (Figure S2), and hence likely holds generally, constituting evidence in favor of a **4POP-IE** explanation of the experimental observations.

### Feasibility of RSG-like oscillatory patterning depends on the ratio of long-range and local transmission delays

Through numerical simulations and bifurcation analyses, we have established above that IP-PING/AP-ING switching can be achieved in both the **4POP-IE** and **4POP-II** models, when the mean cross-hemispheric delay is chosen appropriately. In order to better assess the biological plausibility of each **4POP-XY** motif as a description of RSG dynamics, we sought to understand more precisely how the cross-hemispheric delay interacts with local synaptic kinetics to arrange RSG-like oscillatory patterning. Accordingly, in the Analytical Appendix, we study minimal threshold-linear **4POP** models, which present two key parameters *D*_*i*_ and *D*_*c*_ representing the local inhibitory transmission delay, and the cross-hemispheric transmission delay, respectively. We interpret these model transmission delays as effectively approximating the bulk phase-delay induced by both synaptic filtering and axonal conduction. Hence, in the **QIF-NM** model, the role of *D*_*i*_ is essentially played by the within-hemisphere inhibitory decay time constant *τ*_*I*_ .

For the threshold linear-models, in the regime of small local inhibitory delays (*D*_*i*_ → 0^+^), we show that the coexistence of AP-ING and IP-PING occurs robustly *only for particular ranges of the delay ratio*

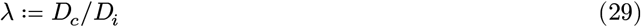

For the **4POP-IE/EI** models, this range is, approximately, 0.36 ≤ *λ* ≤ 2.36, and for the **4POP-II** model, this range is 1.36 ≤ *λ* ≤ 3.36 (Figure S2). This can be considered as an analytical incarnation of the numerical observation, initially depicted in Figure 3B, Figure 4B for the **QIF-NM 4POP** networks, that RSG-like patterning is achieved for shorter cross-hemispheric delays with the **4POP-IE/EI** architecture than with the **4POP-II** architecture.

Although the analytical results are obtained in the asymptotic limit *D*_*i*_ → 0^+^, we further present numerical results in the Analytical Appendix that show a qualitatively similar result holds when *D*_*i*_ takes on physiologically reasonable values, and for the **QIF-NM** models directly (Figure S5, Figure S6). Moreover, we extend our analytical threshold-linear calculations to the **4POP-EE** architecture, in which recurrent excitation is present and cross-hemispheric excitation is the dominant projection. In this case, we show that the appropriate *λ* range is 0 ≤ *λ* ≤ 1.36; thus, to implement RSG-like patterning using the **4POP-EE** architecture would require quite short cross-hemispheric delays.

Short interhemispheric delays, on the same order of magnitude as local inhibitory transmission delays (1-5 milliseconds), are in fact observed across mammalian species ([66]), alongside longer delays on the order of tens of milliseconds ([67]). However, we note again that the majority of superficial RSG pyramidal neurons ([7], [42], [68]) fail to present recurrent E-E connectivity, within or across hemispheres (see Discussion for further detail on this point). Furthermore, although direct long-range inhibitory projections exist in the brain ([64]), this projection type is less common than feedforward long-range inhibition. Hence, current experimental knowledge of long-range projections, together with the presented numerical modeling and analytical results, convergently implicate feedforward **E → I** projections as the more likely contributor to the experimentally observed cross-spectral patterning in RSG.

## Discussion

Extracellular recordings from granular retrosplenial cortex (RSG; [10], [11], [12]) exhibit a remarkable patterning. Band-separated gamma (30-80 Hz) and spline (100-160 Hz) oscillations occur preferentially at distinct phases of the background theta (4-12 Hz) rhythm ([10], [12]), while maintaining divergent interhemispheric phase relationships: gamma bursts are in-phase across hemispheres, while spline bursts are anti-phase ([10]). In this work, we hypothesized that gamma-spline alternation observed *in vivo* is an instance of PING/ING alternation, a previously proposed ([26]) mechanism for cross-frequency coupling in excitatory-inhibitory networks, in which modulation of tonic external drive to inhibitory cells mediates transitions between network regimes producing lower frequency (PING) and higher frequency (ING) oscillations. Indeed, PING/ING alternation has been strongly implicated in the production of similar oscillatory patterning in subiculum ([22]). Thus, the key question we addressed here is to ascertain conditions under which the PING/ING alternation mechanism extends to an inphase PING/anti-phase ING (IP-PING/AP-ING) alternation in interhemispherically coupled excitatory-inhibitory circuits, thereby recapitulating the cross-spectral profile of RSG.

To formalize and tackle this question, we considered networks of neural mass models, assembled into canonical circuit configurations. The **2POP** motif comprises a classic, recurrently coupled excitatory-inhibitory pair of neural masses, while the **4POP-XY** motifs describe symmetrically-coupled excitatory-inhibitory pairs in which interhemispheric communication is dominantly mediated by a single long-range projection type ([45]). When recurrent excitation is weak, as expected in superficial RSG ([41], [42], [11]), **2POP** networks support a unique steady state for any chosen external inputs. Parameter variation (in particular, variation of these external inputs), can subject this steady state to distinct oscillatory instabilities; this is the fundamental mechanism behind PING/ ING alternation ([38], [27], [69]). To next construct IP-PING/AP-ING alternation, we considered instead the unique *symmetric* steady state (s.s.s.) supported by each **4POP** model. This s.s.s. can undergo two types of oscillatory instability: longitudinal, giving rise to oscillations in-phase across hemispheres, and transverse, giving rise to oscillations anti-phase across hemispheres. Thus, our precise goals were 1) to identify the model parameters and associated mechanisms determining whether the s.s.s. approaches a transverse or longitudinal instability, and 2) to identify how the type (transverse/longitudinal) of the instability relates to the frequency of the oscillation generated.

We studied networks constructed from two concrete neural mass models: classical neural activity equations ([70]) with threshold-linear activation function, and next-generation neural mass models which capture exactly the dynamics of globally coupled networks of quadratic integrate-and-fire neurons ([28], [25], [30]). In **4POP** networks built from either model type, we found a key parameter to be the ratio *λ* between long-range and local synaptic delays. Indeed, only for a certain (biologically plausible) range of *λ* values, whose bounds depend on the precise circuit configuration, do the models exhibit a stability landscape consistent with RSG oscillatory patterning. In the Analytical Appendix, we used the threshold-linear **4POP** models to derive an asymptotic approximation for the appropriate *λ* ranges, and determined that the **4POP-IE/EI/II** models all support RSG-like cross-spectral statistics for cross-hemispheric delays reasonably consistent with those previously measured in retrosplenial slice experiments ([42]). Numerical investigation of the next-generation neural mass model also revealed qualitative agreement with these analytical results. This demonstrated, for both models, the existence of broad parameter regimes in which the linearized model dynamics exhibit RSG-like cross-spectral patterning.

We additionally studied the nonlinear dynamics of **4POP** networks tuned to support IP-PING/AP-ING alternation. Here, we used the more realistic next-generation neural mass models, which capture dynamical effects of transient spike synchrony ([25]), and focused attention on the **4POP-IE** and **4POP-II** motifs. We considered that these motifs may constitute informative representatives of two potential circuit scenarios: in the first **4POP-IE** scenario, anti-phase locking of ING oscillations is directly mediated by the excitatory population which also drives local PING oscillations; in the second **4POP-II** scenario, ING oscillations establish anti-phase communication via a pathway independent of the PING-driving excitatory population. Through our numerical explorations, we found that under rhythmic periodic forcing to the inhibitory population (mimicking theta-rhythmic GABAergic inputs onto RSG I-cells from medial septum; [71]), both **4POP-IE** and **4POP-II** models exhibited IP-PING/AP-ING switching solutions. In the latter model, but not the former, these switching solutions were chaotic, reflective of underlying bistability between IP-PING and AP-ING limit cycles in the unforced **4POP-II** network. Furthermore, we found that symmetry-breaking bifurcations of the s.s.s. occur in distinct parameter regimes for the **4POP-IE** and **4POP-II** models, such that transient symmetry-breaking events occurring during theta-rhythmic states are coupled to distinct phases of the theta rhythm. Lastly, in the presence of multiplicative noise mimicking finite-size effects, both the **4POP-IE** and **4POP-II** models produce cross-spectra similar to bilateral RSG recordings, with **4POP-IE** exhibiting the clearest agreement with experimental metrics.

Thus, our work establishes that RSG-like oscillatory patterning can robustly occur in simple, biophysically motivated, models of minimal interhemispheric circuitry. We have accomplished this by providing a detailed description of the stability landscape for **4POP** symmetrically coupled E-I pairs, contributing to the existing literature focused on PING-ING alternation in **2POP** pairs ([38], [27], [69]) and the literature which studies phase-locking of oscillations in symmetric delay-coupled neural mass networks ([72], [73], [74], [75], [76]). Nonetheless, we note several limitations of our work which provide important targets for future computational/mathematical studies. First, since our networks are comprised of QIF neurons, they are an exact description only for networks consisting of Type I ([77], [78]) neurons. While prior studies have examined dependence of neuronal excitability type on PING/ING dynamics ([27]), it will be important to understand how the choice of excitability type affects dynamics in interhemispheric circuits of the kind studied here. Furthermore, gap junctions are a ubiquitous feature of cortical inhibitory networks ([79], [80]); hence, it will be important to understand how the presence of gap junctions affects the bifurcation structures examined here. Fortunately, gap junctions are easily incorporated into the **QIF-NM** framework ([31], [81])

To conclude, we interpret the results in the context of extant experimental data and emphasize several key takeaways. We claimed that cross-hemispheric inhibition, mediated by feedforward **E → I** projections, may be crucial for establishing the cross-spectral profile exhibited by RSG. This is based in the assumption that recurrent excitation is weak within and across hemispheres, which is indeed true for the dominant excitatory cell type in superficial RSG, the low-rheobase (LR) cell ([41], [7], [42]). However, LR cells reside alongside a smaller ([68]) population of superficial regular spiking (RS) neurons, which do exhibit recurrent excitation. It is therefore natural to ask whether the RS-inhibitory subcircuit may implement the observed cross-spectral patterning via the **4POP-EE** architecture. We note that existing *in vitro* experimental results indicate cross-hemispheric inputs to LR and RS cells exhibit comparable latencies, around 7-12 milliseconds ([42]). Although measurements of inhibitory synaptic decay times are not available for RSG, we cite past experiments reporting values on the order of 1-3 milliseconds in the hippocampal formation ([82]). Thus, we argue that whether considering the RS-inhibitory or LR-inhibitory subcircuit as the primary rhythm generator, these values are indeed more compatible with the *λ* ranges presented for **4POP-IE/II** than for **4POP-EE**. Furthermore, our analytical results were consistent with the numerical observations of closer PING and ING frequencies in the **4POP-II** model than in the **4POP-IE** model. These analyses suggest this qualitative feature may exhibit weak dependence on the choice of model or parameters, hence providing further evidence in favor of the **4POP-IE** motif over the **4POP-II** motif to explain the experimental findings. Nonetheless, rigorously testing this hypothesis will require careful future experiments, ideally synthesizing *in vitro* measurements with optogenetic manipulation of cell-type specific callosal projections ([7]).

Regardless of network architecture, requiring RSG-like cross-spectra constrains the allowable range of cross-hemispheric delays, suggesting that experimentally observed phase-relationships is likely to depend on well-tuned axonal delays. We emphasize that potentiation of synapses with specific delays can be achieved in a self-organized manner, through biologically plausible plasticity rules ([83]), and it will be interesting for future work to ascertain whether RSG-like cross-spectral patterning is a stable configuration for plastic interhemispheric networks with delays. Moreover, we argue that since delays are so crucial to this patterning, disrupted RSG phase relationships could be expected in demyelinating diseases, such as multiple sclerosis, or during early life ([84]). Hence, we additionally suggest as fruitful experimental directions, the exploration of macroscopic RSG dynamics in model organisms for such diseases, as well as during the neurodevelopmental trajectory.

## Acknowledgements

This work was initiated during the ICERM 2023 Math+Neuroscience workshop; we thank Peter Thomas, Amit Bose, Saafan Sadiq, and Max Kreider for helpful conversations. This work was supported by NIH T32DA060142; NIH R34NS127101; NIH P50NS123067; NIH R01MH129282; Alzheimer’s Association Grant AARG-NTF-21-846572;

Berger Endowment, G102865/321033.

## Statement on use of AI tools

The first author used a large language model to facilitate proofreading and formatting of the manuscript text. Models, analyses, simulations, and figures were developed without use of AI tools. The authors have reviewed all edits and take full responsibility for the content of this work.

## Addendum 1: Existence and uniqueness of 2POP steady state

In the **2POP** model including recurrent excitation, the steady states satisfy

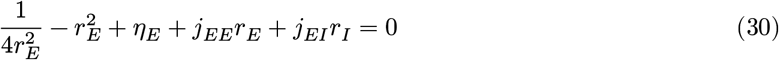

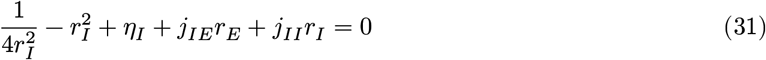

Assume the biological sign constraints *j*_*II*_, *j*_*EI*_ < 0 and *j*_*EE*_, *j*_*IE*_ > 0. We have *r*_*I*_ = *f*(*r*_*E*_) where

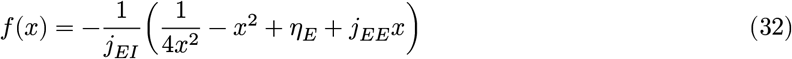

which we will show is strictly decreasing for sufficiently small *j*_*EE*_. However, implicitly differentiating the second equation shows *r*_*I*_ = *g*(*r*_*E*_) where *g* is positive and strictly increasing; specifically, we find

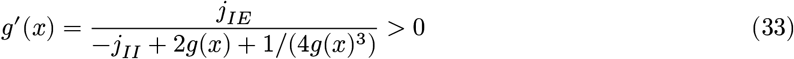

Thus, under these conditions there exists a unique steady state. Finally, for *f* to be strictly decreasing, we need *f* ^′^(*x*) < 0, i.e. the function *x* ↦ −1/(2*x*^3^) − 2*x* + *j*_*EE*_ should have a negative maximum; this happens precisely when *j*_*EE*_ < (8/3) ⋅ (3/4)^1/4^ ≈ 2.4816. In particular, for *j*_*EE*_ = 0 as considered in this work, the **2POP** model admits a unique steady state, and the **4POP** models admit a unique symmetric steady state.

## Addendum 2: Equivalence of local bifurcations for 4POP-IE and 4POP-II

For the **4POP-IE/EI** models, we have

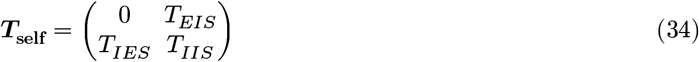

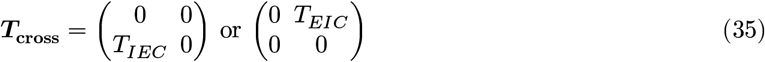

depending on the model. For fixed ***j***_tot_, the steady state rates are equal across both models. The bifurcation loci satisfy 0 = det(***I*** − ***T***_self_ − *ε****T***_cross_) where *ε* = ±1 determines longitudinal/transverse bifurcations, respectively. For **4POP-IE**, the determinant evaluates to

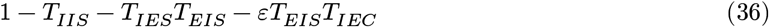

and for **4POP-EI**, it evaluates to

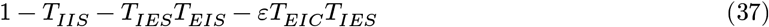

Note these expressions are identical, except for the last term in each case; we have

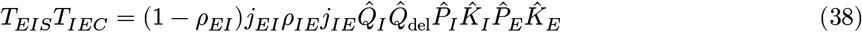

and

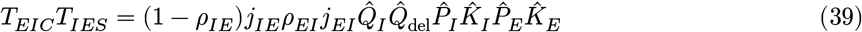

where 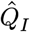 is the first-order inhibitory synaptic filter and 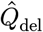 is the interhemispheric delay filter. It is now clear that these expressions become identical if *ρ*_*EI*_ = *ρ*_*IE*_.

## Analytical Appendix

### Linear response properties of neural mass networks

#### Network equations

Consider a general network of 1 ≤ *α* ≤ *P* neural mass models (NMMs) with current-based coupling, each node representing the spatially averaged dynamics of a neuronal population:

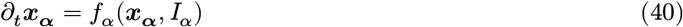

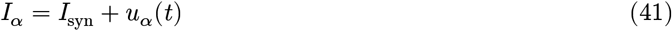

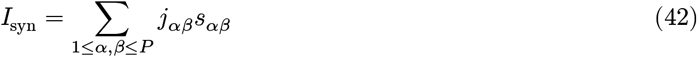

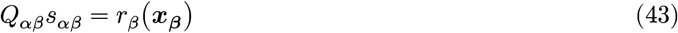

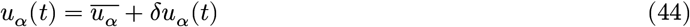

Here ***x***_***α***_ is the latent dynamical state associated with node *α*, taking values in some Euclidean space 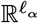.

Nodes interact through their *firing rates*, specified as an observable 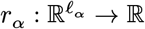. The synapse from node *β* to node *α* has associated synaptic variable *s*_*αβ*_, related with the firing rate *r*_*β*_(***x***_***β***_) through some linear differential operator *Q*_*αβ*_. We assume throughout that *Q*_*αβ*_ is always normalized, with impulse response integrating to one.

Lastly, each input current *I*_*α*_ into node *α* is comprised of: **1)** a recurrent network component *I*_syn_, obtained as the weighted sum of synaptic variables *s*_*αβ*_ with coupling strengths *j*_*αβ*_, and **2)** an external input *u*_*α*_(*t*) given as the sum of a DC-component 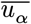 which is constant in time, and fast-varying low-amplitude AC-component *δu*_*α*_(*t*).

To interpret the setup within the context of this work, we consider that the DC-input 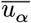 models a slowly-varying theta-rhythmic input to the RSG network, while fast noise inputs *δu*_*α*_ induce, through network filtering, the high-frequency oscillations observed on the LFP.

#### Network linearization

In the absence of fast fluctuations *δu*_*α*_, the network dynamics are deterministic and we assume the system settles to a stable equilibrium 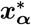. We denote the steady state firing rates 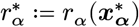 and steady state inputs 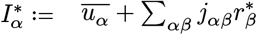.

We linearize about this steady state to approximate the dynamics with fast fluctuations included. Writing

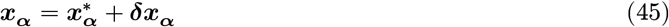

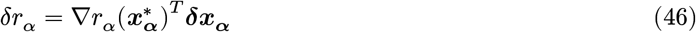

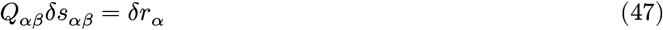

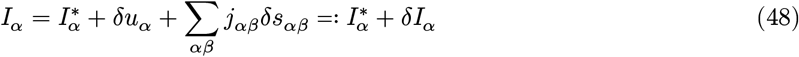

we then have the tangent dynamics

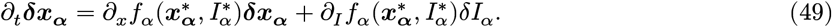

Next, we take the Laplace transform to obtain

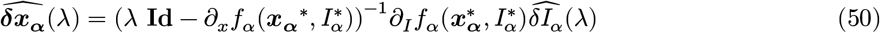

where *λ ∈* ℂ and setting *λ* = *iω* for *ω ∈* ℝ recovers the Fourier transform. Accordingly, the firing rate fluctuations are obtained as 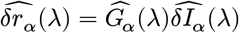 where we define the quantity

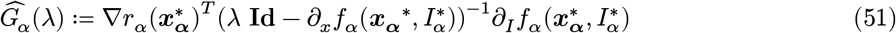

as the *intrinsic transfer function* of the population *α*. This quantity can be understood as encoding how microscopic dynamical features (e.g. adaptation, gap junctions, etc.) of the individual neurons comprising population *α* affect the macroscopic linear response properties of the population as a whole. It follows that

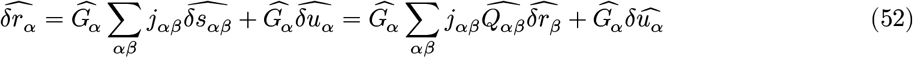

where we define 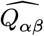 as the Laplace transform of the impulse response for *Q*_*αβ*_. Thus, stacking these quantities into vectors ***δr, δu*** *∈* ℝ^*P*^, we have 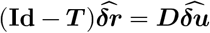, where the *network transfer matrix* ***T*** is defined by 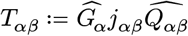 and ***D*** denotes the diagonal matrix with 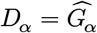. Accordingly, the characteristic equation of the network is det(**Id** − ***T*** ) = 0, and the cross-spectrum upon providing inputs *δu*_*α*_ is

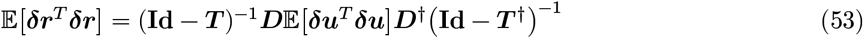

where the expectation in the middle is the input noise covariance.

#### Example neural mass models

In this appendix, we will study two canonical neural mass models. In the *Wilson-Cowan-style rate equations* (model **WC-NM**), the state ***x***_***α***_ is one-dimensional; we call this scalar state variable *R*_*α*_ and take

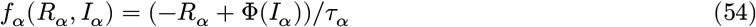

for some positive, monotone increasing nonlinear function **Φ** and population time constant *τ*_*α*_. For concreteness, we consider the family

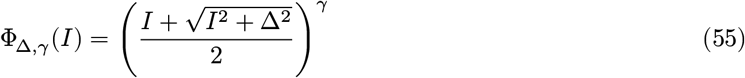

where Δ represents the smoothing by microscopic noise and *γ* is the population output nonlinearity. In the *quadratic integrate-and-fire neural mass* (model **QIF-NM**), the state ***x***_***α***_ is two-dimensional; we call the state variables *R*_*α*_ and *V*_*α*_, and take

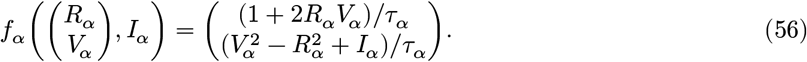

In both cases, to prescribe the firing rate output, we simply set *r*_*α*_ = *R*_*α*_. Note that the steady state firing rate of the QIF-NM model, as a function of the input *I*_*α*_, is just **Φ**_1,1/2_, and hence embeds directly into the family **Φ**_Δ,*γ*_ . Indeed, networks of the **WC-NM** type can be obtained directly as a limiting case of **QIF-NM** networks, when the synaptic kinetics are made arbitrarily slow ([85]).

We proceed to compute the intrinsic transfer function for each model, at some arbitrary network steady state having rates 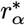. For the family **Φ**, the population DC-gain (i.e., 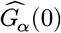) can be written explicitly as a function of the output rate:

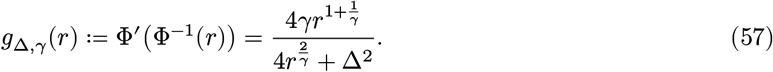

In particular, we have *g*_1,1/2_(*r*) = 2*r*^3^/(4*r*^4^ + 1); it is convenient to define also define *h*_*k*_(*r*) = 2*r*^*k*^/(4*r*^4^ + 1) so that *g*_1,1/2_ = *h*_3_. Then, for the **WC-NM**, we obtain

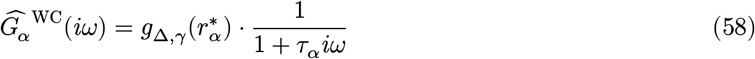

This is a first-order low pass filter, for all 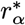.

In contrast, for the **QIF-NM**, one obtains

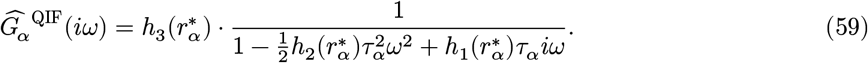

This is a second-order filter, which behaves as low-pass for 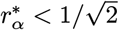, and for 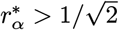 has a resonant peak at 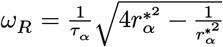. Note that, by construction, both intrinsic transfer functions have the same DC-gain: 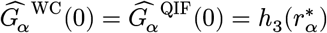. However, in the **WC-NM** model, effects of output rate 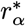 and frequency *ω* are separable, while for the **QIF-NM**, the rate parameter 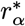 itself induces phase effects.

### Excitatory-inhibitory module

Consider a two-population excitatory inhibitory module, corresponding to the **2POP** motif described in the main text; we use subscripts *e, i* to denote the excitatory and inhibitory populations, respectively. One then has the characteristic equation

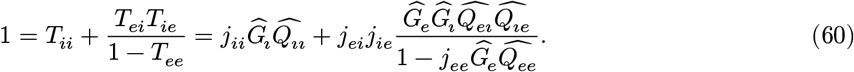

Previous work ([38]) has extensively described the linear instability scenarios for this E-I module. In particular, Hopf instabilities can arise in two qualitatively distinct ways: the excitatory-inhibitory feedback PING mechanism, or the inhibitory-inhibitory feedback ING mechanism. Accordingly, we let *α*_*I*_ = *j*_*ii*_ ≤ 0, *α*_*P*_ = *j*_*ei*_*j*_*ie*_ ≤ 0 be parameters which quantify the tendency of the network oscillation to rely on one or the other qualitative mechanism. Further, let *Q ≔* {*α*_*I*_ ≤ 0, *α*_*P*_ ≤ 0} denote the quadrant of the parameter plane corresponding to biologically relevant weights. Defining 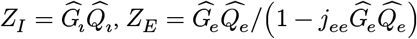, and *Z*_P_ = *Z*_*I*_*Z*_*E*_, the characteristic equation is simply

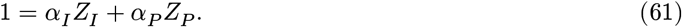

We consider the Hopf bifurcation locus in the (*α*_*I*_, *α*_*P*_ ) parameter plane; since *α*_*I*_, *α*_*P*_ are real-valued weights, one obtains a convenient parametric representation of the main Hopf branch by solving the linear system:

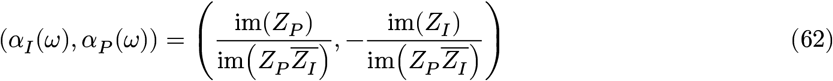

which is valid for any *ω* > 0 such that 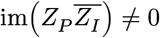. Additionally, considering *ω* = 0, one obtains the fold locus given by the line

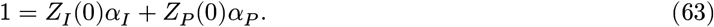

Note that this fold line intersects *Q* only for *Z*_*P*_ (0) < 0, yielding the inhibition-stabilized regime (ISN; [44]). The fold locus and Hopf curve meet at the Bodganov-Takens point 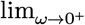*α*_*I*_(*ω*), *α*_*P*_ (*ω*)).

#### Symmetrically coupled E-I modules

As in the main text, we consider a pair of symmetrically coupled E-I modules, with full synaptic weight matrix taking the ℤ_2_-symmetric form

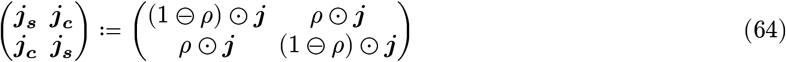

where

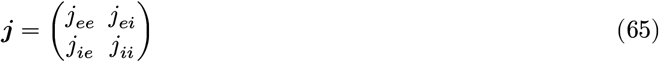

describes the total coupling strength and

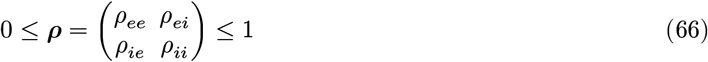

are the interhemispheric projection ratios. Further, recall the rank-1 network motifs considered in the main text, namely the **4POP-XY** architectures; for a connection type *xy*, this architecture comprises networks for which *ρ*_*xy*_ is the only nonzero entry of ***ρ***.

Each four population network admits distinct *longitudinal* and *transverse* Hopf instabilities, respectively giving rise to in-phase and anti-phase oscillations. In both these cases, the corresponding characteristic equations take precisely the same form as for the single E-I module, except with effective synaptic filters which present a linear combination of the local and long-range filtering. Precisely, denote by 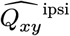 the local synaptic filter and 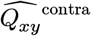 the long range synaptic filter, for each connection type *xy*. The overall effective filter on connection type *xy* is

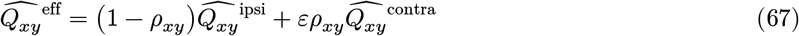

where *ε* = ±1 gives the expression for longitudinal/transverse bifurcations, respectively.

In particular, the longitudinal/transverse Hopf curves are again obtained parametrically. To fix notation for what follows, let ***α***_**2**_(*ω*) denote the Hopf curve for the **2POP** network, let 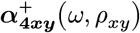 denote the longitudinal Hopf curve for the **4POP-XY** network, and let 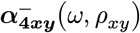 denote the transverse Hopf curve for the **4POP-XY** network. Note that

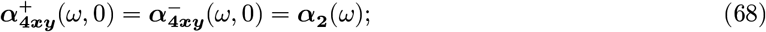

the **4POP** curves degenerate to the **2POP** curve when interhemispheric projections vanish. For *ρ*_*xy*_ > 0, the curves 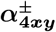 are perturbed from ***α***_**2**_ and intersect at a sequence of double Hopf points. Let (*ω*_+_, *ω*_−_) be such a double Hopf point, i.e. a solution to 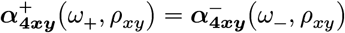. Suppose there exists some *ω*_0_ such that as *ρ* → 0^+^ we have

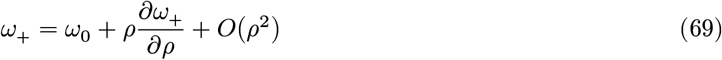

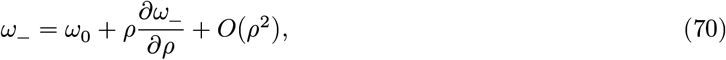

where the derivatives are evaluated at *ρ* = 0. Then, solving the defining equation for the double Hopf point, at first order in *ρ*, gives the necessary condition

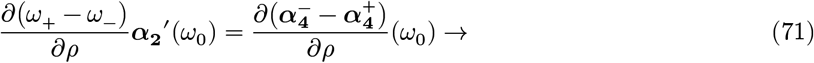

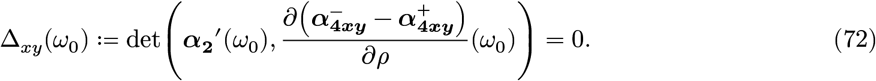

Hence, in the weak coupling limit *ρ*_*xy*_ → 0^+^, the double Hopf points are in direct correspondence with the zeros of Δ_*xy*_. Solutions *ω*_0_ are critical frequencies at which the network oscillation switches from in-phase to anti-phase, or vice versa.

### Explicit example: threshold-linear WC-NM network

We work out an example analytically, namely the classical rate E-I module with Dirac delta delay kernels:

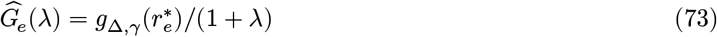

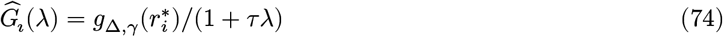

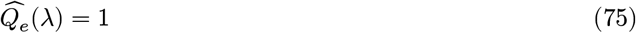

and

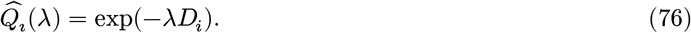

This is the extreme instance of the *D*_*e*_ ≪ *D*_*i*_ regime considered by [38], in which the transition between PING and ING oscillations occurs continuously, along a single Hopf branch. Here, we have set simply *D*_*e*_ = 0.

Importantly, in what follows, we will make an additional assumption which renders the calculations more tractable: we take Δ = 0, *γ* = 1; this *threshold linear* assumption yields 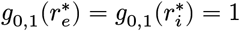, so that the local bifurcation structure is totally independent of the external inputs 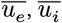 However, in the limiting case *j*_*ee*_ → 0, the separable structure of the intrinsic transfer function for **WC-NM** networks can be exploited to show how these bifurcations induced by the network coupling structure translate to bifurcations in the 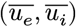-space, even when Δ > 0. We will demonstrate this later on, after studying the stability landscape for the **2POP** and **4POP** threshold-linear networks.

#### Threshold-linear 2POP stability landscape

Throughout, we assume Δ = 1, *γ* = 0, so that we are examining the same threshold-linear model studied in [38]. We obtain the Hopf curve

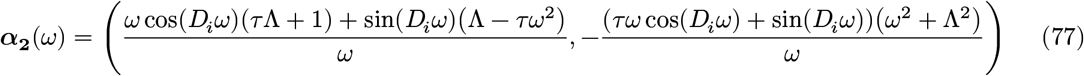

and fold line

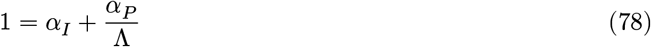

where Λ *≔* 1 − *j*_*ee*_. As *ω* → 0^+^, we have

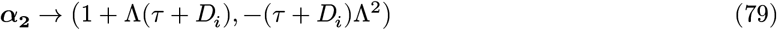

which locates the Bogdanov-Takens point. In a non-ISN, this point is outside *Q*; in fact, it enters *Q* only once *j*_*ee*_ > 1 + 1/(*τ* + *D*_*i*_), at which point both feedback inhibition (as quantified by *α*_*P*_ ) and recurrent inhibition (as quantified by *α*_*I*_ ) are required to stabilize the network.

**Figure S1:**
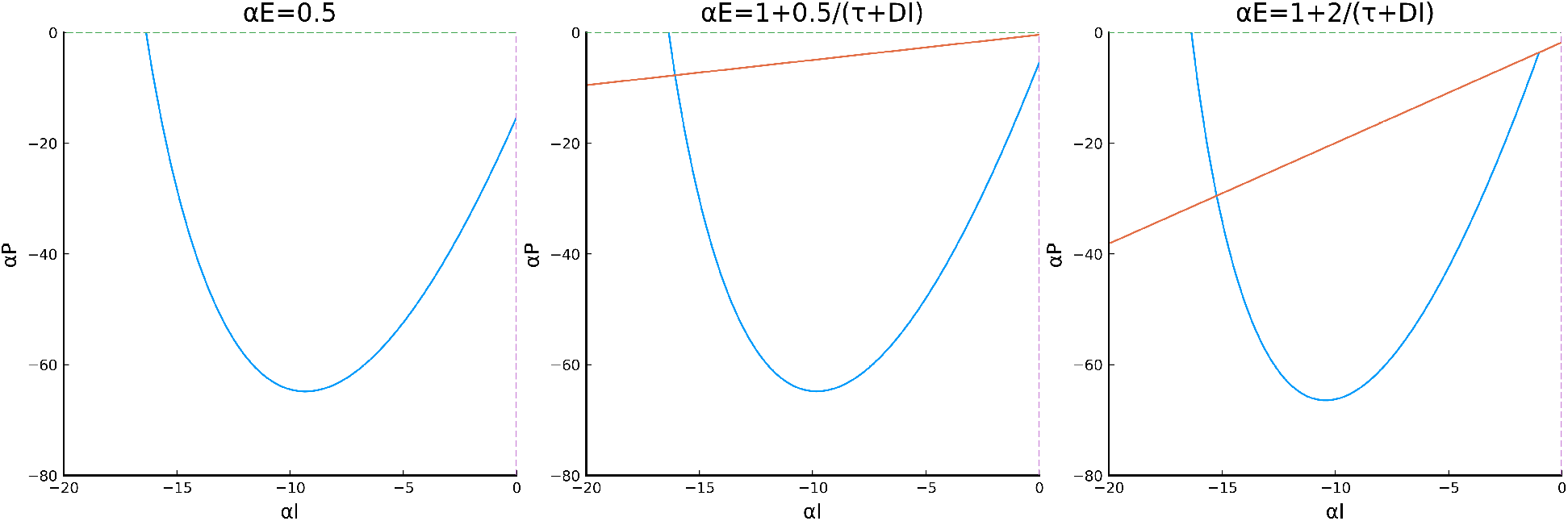
Bifurcation curves for threshold-linear **2POP** when varying *α*_*E*_. Blue = Hopf, red = fold. Parameters *D*_*i*_ = 0.1, *τ* = 1.

The pure ING instability occurs when *α*_*P*_ = 0, at the frequency *ω*_*I*_ given by the smallest positive solution to

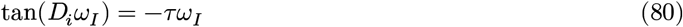

while the pure PING instability occurs when *α*_*I*_ = 0, at the smallest positive solution *ω*_*P*_ to

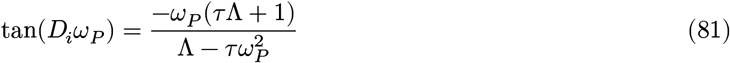

Thus, the portion of the ***α***_**2**_ curve in *Q* is swept out over the frequency range [*ω*_*P*_, *ω*_*I*_ ]. As *D*_*i*_ → 0^+^, we have the asymptotics *ω*_*I*_ ∼ π/(2*D*_*i*_) and 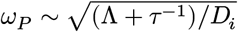. In particular, the frequencies diverge as the delay vanishes.

A natural question is to ask how the 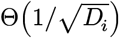 frequency scaling in the PING regime transitions into Θ(1/*D*_*i*_) frequency scaling in the ING regime. To address this, we set *v* = *α*_*P*_ /*α*_*I*_, which trace out the Hopf line in *Q* when *v* ranges over [0, ∞], (as long as we are in the non-ISN, *j*_*ee*_ < 1). Then, the frequency *ω*(*v*) of the Hopf point at slope *v* satisfies

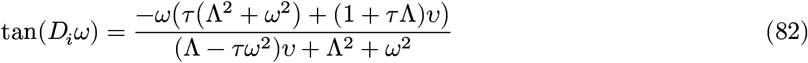

so that *ω*(0) = *ω*_*I*_ and *ω*(∞) = *ω*_*P*_ as above. Assume *v* = *ξ*/*D*_*i*_ as *D*_*i*_ → 0^+^, for some constant *ξ ∈* (0, ∞). Then via dominant balance arguments, one obtains

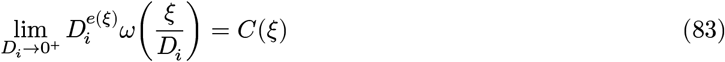

where

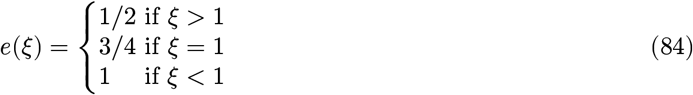

and

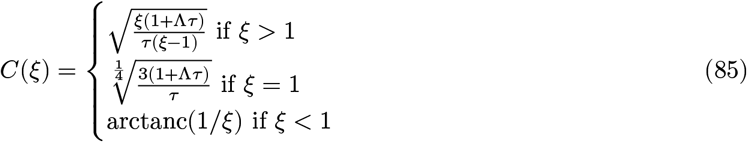

where arctanc(*x*) is the unique *w ∈* (0, π/2) which satisfies tan(*w*)/*w ≕* tanc(*w*) = *x*.

#### Counting threshold-linear 4POP double Hopf points

To specify the threshold-linear **4POP** models fully, we must describe the choice of synaptic kinetics across and within hemispheres. We consider 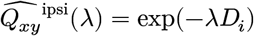 if *y* = *i* (presynaptic inhibitory) and 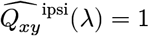 if *y* = *e* (presynaptic excitatory); similarly, we set 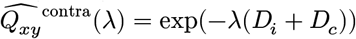 for *y* = *i* and 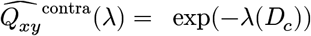 for *y* = *e*. Then, for each **4POP-XY** model, we have

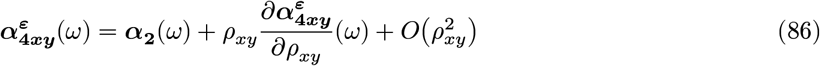

with the partial derivative evaluated at *ρ*_*xy*_ = 0 and *ε* = ±. A tedious calculation gives

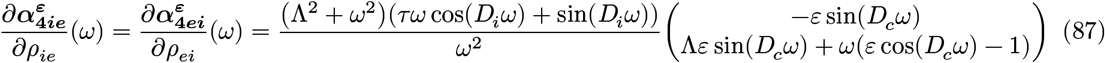

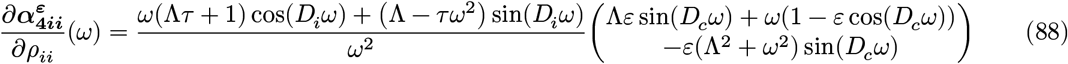

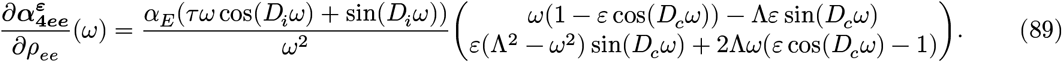

Now, considering the limit as *D*_*i*_ → 0^+^ and using that *ωD*_*i*_ = *O*(1) in this limit, we write *θ ≔ ωD*_*i*_ and *λ ≔ D*_*c*_/*D*_*i*_. Then, another tedious calculation using the above yields

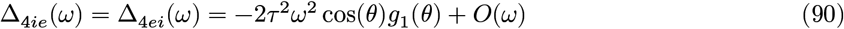

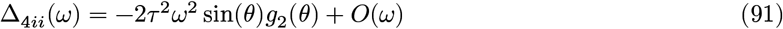

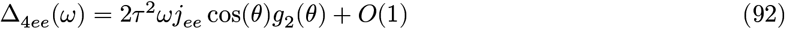

where

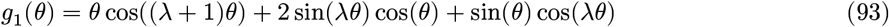

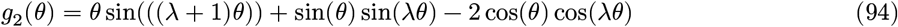

Thus, in the limit *D*_*i*_ → 0^+^, nontrivial (i.e., not *θ* = 0 or *θ* = π/2) double Hopf points corresponding with intersections of 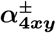 are in bijection with roots of *g*_1_ for the **4POP-IE/EI** models and in bijection with roots of *g*_2_ for the **4POP-II/EE** models. Moreover, we observe that sign(Δ_4*ii*_) = −sign(Δ_4*ee*_), indicating that while double Hopf points occur at the same critical frequencies for both **4POP-II** and **4POP-EE**, the Hopf intervals between double Hopf points are of opposite sign (Figure S2), i.e. if **4POP-EE** admits longitudinal instability at a certain frequency-delay pairing, then **4POP-II** would admit transverse instability at the same pairing.

**Figure S2:**
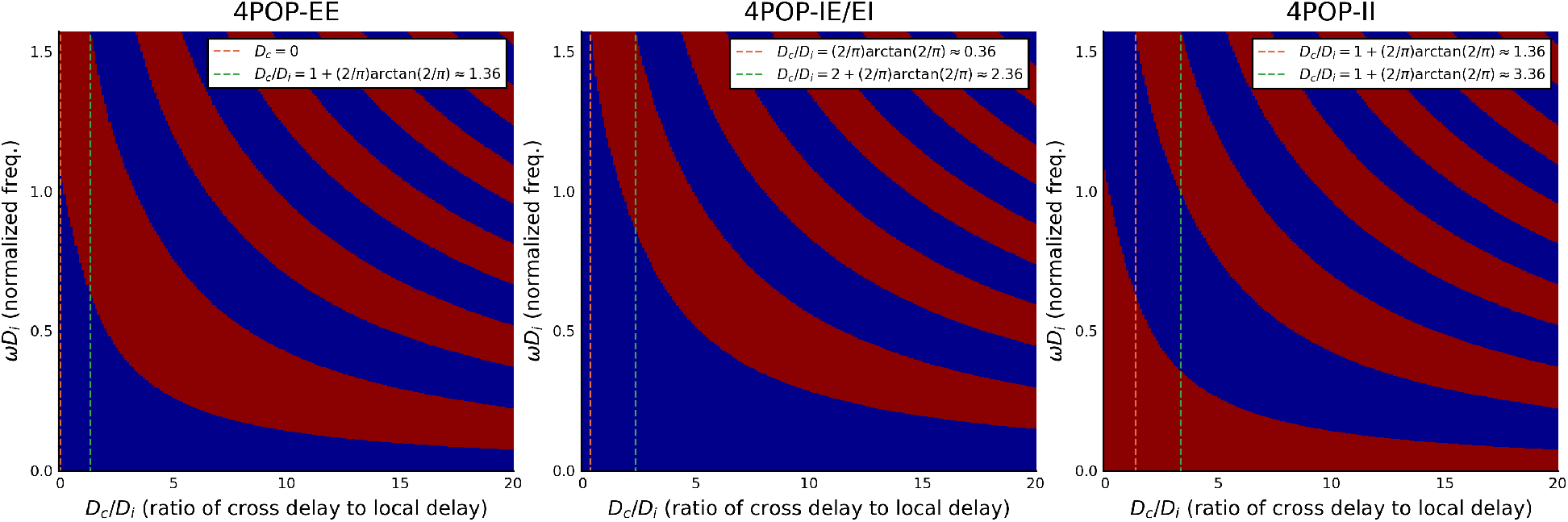
Hopf bifurcation type for **4POP** threshold-linear models, in the limit *D*_*i*_ → 0^+^. Blue = longitudinal, red = transverse.

Of crucial interest for this work is to identify the conditions under which anti-phase high-frequency oscillations (which rely more heavily on the ING pathway; AP-HF) can coexist with in-phase lower-frequency oscillations (which rely more heavily on the PING pathway; IP-LF). In the case of large *λ* = *D*_*c*_/*D*_*i*_ there are two relevant biological considerations. First, although large *λ* induces a proliferation of double Hopf points, the interleaving Hopf intervals are shorter. Thus, even if the network is poised near a double Hopf point with nearby high-frequency anti-phase and low-frequency in-phase oscillations, the achievable frequency ranges are shorter (before moving far enough to trigger an interhemispheric phase shift inconsistent with the retrosplenial data). Second, the assumption of a Dirac delay kernel is unrealistic, and a more realistic kernel accounting for distributed delays (e.g. a Gamma kernel of the type considered in the main text) is likely to have a saturating phase response; as observed in our numerical simulations, this seems to mitigate stable phase-locking for large cross-hemispheric delays.

Thus, we consider that the most plausible scenarios are those for which *D*_*c*_ is a relatively small multiple of *D*_*i*_. Accordingly, for each **4POP** motif, we compute the smallest interval of *λ* values in which anti-phase pure ING coexists with in-phase lower-frequency oscillations. Letting *C* = (2/π) arctan(2/π), for **4POP-EE**, the relevant range is 0 ≤ *λ* ≤ 1 + *C* ≈ 1.36, indicating that E-E cross-hemispheric projections can yield a network which produces AP-HF/IP-LF even in absence of explicit cross-hemispheric delays. Realistically, however, such delays are likely to exist, suggesting that producing the experimentally observed patterning through E-E connections alone is implausible. For **4POP-IE/EI**, the interval is 0.36 ≈ *C* ≤ *λ* ≤ 2 + *C* ≈ 2.36; and for **4POP-II**, the interval is 1.36 ≈ 1 + *C* ≤ *λ* ≤ 3 + *C* ≈ 3.36. These ranges contain more realistic values for *λ*. However, we note that in the case of **4POP-II**, the AP-HF/IP-LF oscillations do not comprise the entire frequency range achievable by the network, coexisting with an additional AP-LF range contiguous to the pure PING at frequency 0 (Figure S2, right panel). This is consistent with the observation of closer IP-PING/AP-ING frequencies for **4POP-II** in our numerical simulations of the QIF-NM model.

#### Threshold-linear 4POP fold instabilities

We study the fold lines obtained analytically for the **4POP** threshold-linear networks:

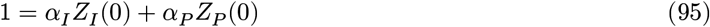

where 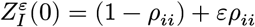 and

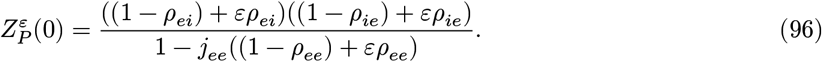

When *ε* = 1, this gives 1 = *α*_*I*_ + *α*_*P*_ /(1 − *j*_*ee*_), recovering the **2POP** fold line (Equation 63) for longitudinal instabilities. When *ε* = −1, we obtain the transverse fold line

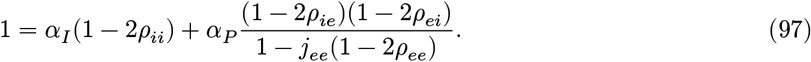

which has *α*_*P*_ -intercept

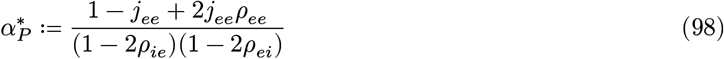

and slope 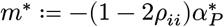. In the non-ISN regime *j*_ee_ < 1, for sufficiently small ***ρ*** we have 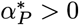 and *m*^*^ < 0, so the line misses *Q* entirely. It can be made to intersect *Q* by:

- increasing *ρ*_*ii*_ beyond 1/2 to flip the sign of *m*^*^. In this case, since 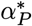 is unchanged, the intersection of the line with the *α*_*I*_-axis comes in from *α*_*I*_ = −∞. This is the **4POP-II** situation described in the main text.
- increasing *ρ*_*ei*_ or *ρ*_*ie*_ beyond 1/2 to flip the sign of 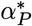. This forces *m*^*^ > 0 (as long as *ρ*_*ii*_ remains less than 1/2), and the intersection of the line with the *α*_*P*_ -axis comes in from *α*_*P*_ = −∞. This is the **4POP-IE/EI** situation described in the main text.

In the ISN regime *j*_*ee*_ > 1, taking 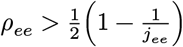 compensates the sign of the numerator of 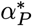, so that the same behaviors hold when *ρ*_*ie*_, *ρ*_*ei*_, or *ρ*_*ii*_ are modulated. However, if each hemisphere is locally inhibition stabilized, but *ρ*_*ee*_ is too small, then transverse instability will occur (i.e., the transverse fold line intersects *Q*) unless *ρ*_*ie*_ or *ρ*_*ei*_ is chosen large enough (> 1/2) to compensate; in this case, increasing *ρ*_*ie*_ or *ρ*_*ei*_ up to 1/2 sends 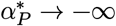, and increasing beyond 1/2 makes 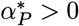. We note that interhemispheric I-I projections cannot mitigate this transverse instability, since 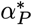 is independent of *ρ*_*ii*_. This analysis is for the threshold linear **WC-NM 4POP** networks, but analogous results are seen to hold for **WC-NM** modules with nonlinear population response, as well as for the **QIF-NM** module.

### Input-space bifurcations for the E-I module with *j*_*ee*_ = 0

We now return to the question of how the coupling strength bifurcations studied above can yield insight into input space bifurcations, which are more directly relevant to capture the PING-ING alternation mechanism examined in the main text. Consider the **2POP WC-NM** network with *γ* = 1, but Δ > 0, and no recurrent excitation *j*_*ee*_ = 0. However, we will again consider that there is an inhibitory delay, 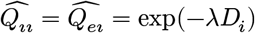 and 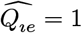. In this case, the characteristic equation is

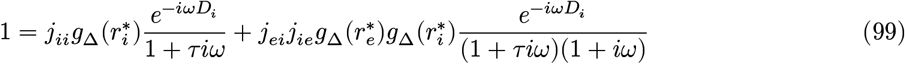

Just as in the calculations for the threshold-linear case, one can solve a two-dimensional linear system to obtain

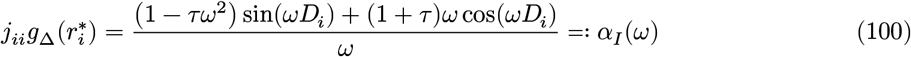

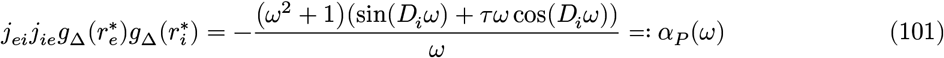

Note that in the chosen scenario, we have *g*_Δ_(*r*) = 4*r*^2^/(4*r*^2^ + Δ^2^), which is a sigmoidal map with image (0, 1); its inverse is 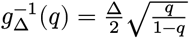 for *q ∈* (0, 1). We find that for a particular frequency *ω ∈* [*ω, ω* ], there is a solution in rates 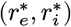 if and only if |*α*_I_ (*ω*)| < |*j*_*ii*_| and |*α*_P_ (*ω*)/*α*_I_ (*ω*)| < |*j*_*ei*_ *j*_*ei*_ /*j*_*ii*_|. Thus, for fixed coupling weights, we find that any amount of microscopic noise Δ fully precludes expression of all the lowest frequencies in [*ω*_*P*_, *ω*_*I*_ ], since the slope *v* = *α*_*P*_ /*α*_*I*_ is bounded above.

Letting *ω*_*M*_ be the smallest positive frequency such that *v*(*ω*) = *j*_*ei*_*j*_*ie*_/*j*_*ii*_, we find that the Hopf curve in the 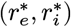-plane is traced by frequencies (*ω*_*M*_, *ω*_*I*_]. For simplicity, assume |*j*_*ii*_| > |*α*_*I*_ (*ω*_*I*_)| to guarantee existence of 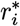. Then, as 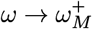, we see that 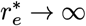 while 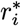 approaches the value 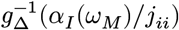. As 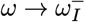, we have 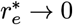 and again 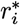 approaches the value 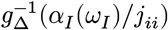.

Thus, for a fixed set of coupling weights, we have computed analytically a curve of Hopf points in the 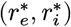 plane capturing the PING-ING transition. Points on this curve have 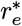 varying from 0 to ∞ while 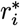 varies from its maximum 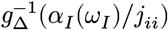 to its minimum 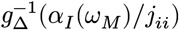. In particular, this demonstrates proximity of stable steady states with high 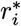 and low 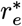 to a high-frequency ING-like Hopf instability, while stable steady states with high 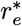 and low 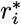 are closer to a low-frequency PING-like Hopf instability.

A final translation into the input space 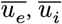 is straightforward because the model structure is such that for fixed coupling weights, there is a unique set 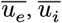 achieving any given 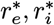, namely

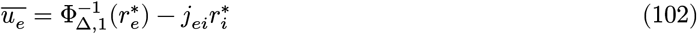

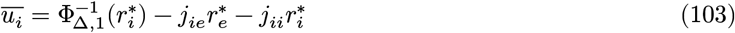

where

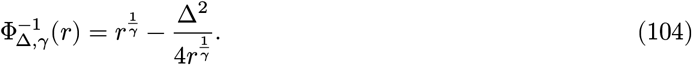

**Figure S3:**
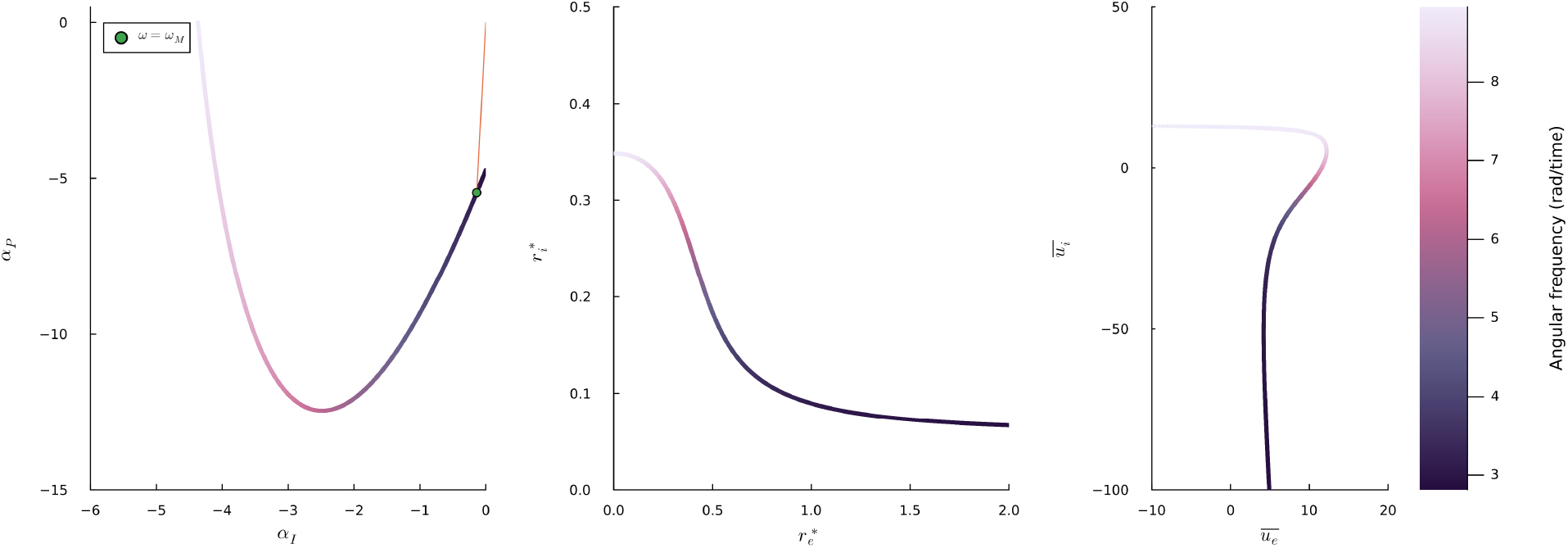
Weight-space Hopf bifurcation curve for **2POP WC-NM** with *γ* = 1, translated into input space. Here we have set *j*_*ee*_ = 0, *j*_*ei*_ = −40, *j*_*ie*_ = 40, *j*_*ii*_ = −40, Δ = 1, *τ* = 0.5, *D*_*i*_ = 0.2. Marker in leftmost plot shows *ω*_*P*_, the infimum of all frequencies for which there exists some 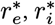 satisfying the Hopf characteristic equation.

In Figure S3, we show how for *γ* = 1, the **WC-NM** Hopf curve in weight space corresponds with a unique Hopf curve in input space. Similarly to the curve depicted in Figure 2 of the main text for the **QIF-NM** model, there is a range of 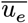 for which varying 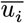 parallel to the Hopf curve can transition between high- and low-frequency Hopf points. For *γ* > 1, the picture is similar, since *g*_Δ_ is then increasing on its domain and again there is at most one Hopf point in rate space for each Hopf point in weight space. In contrast, for *γ* < 1, *g*_Δ_ is unimodal, and each Hopf point in weight space consequently yields four corresponding points in rate space; this more complex bifurcation structure can be considered as somehow more analogous to the classical Wilson-Cowan model with sigmoidal firing rate functions ([86]). Nonetheless, all these **WC-NM** cases are tractable from separability of the intrinsic transfer function; in the **QIF-NM** case, the rate-dependent phase effects complicate the analysis further and preclude using the same technique.

**Figure S4:**
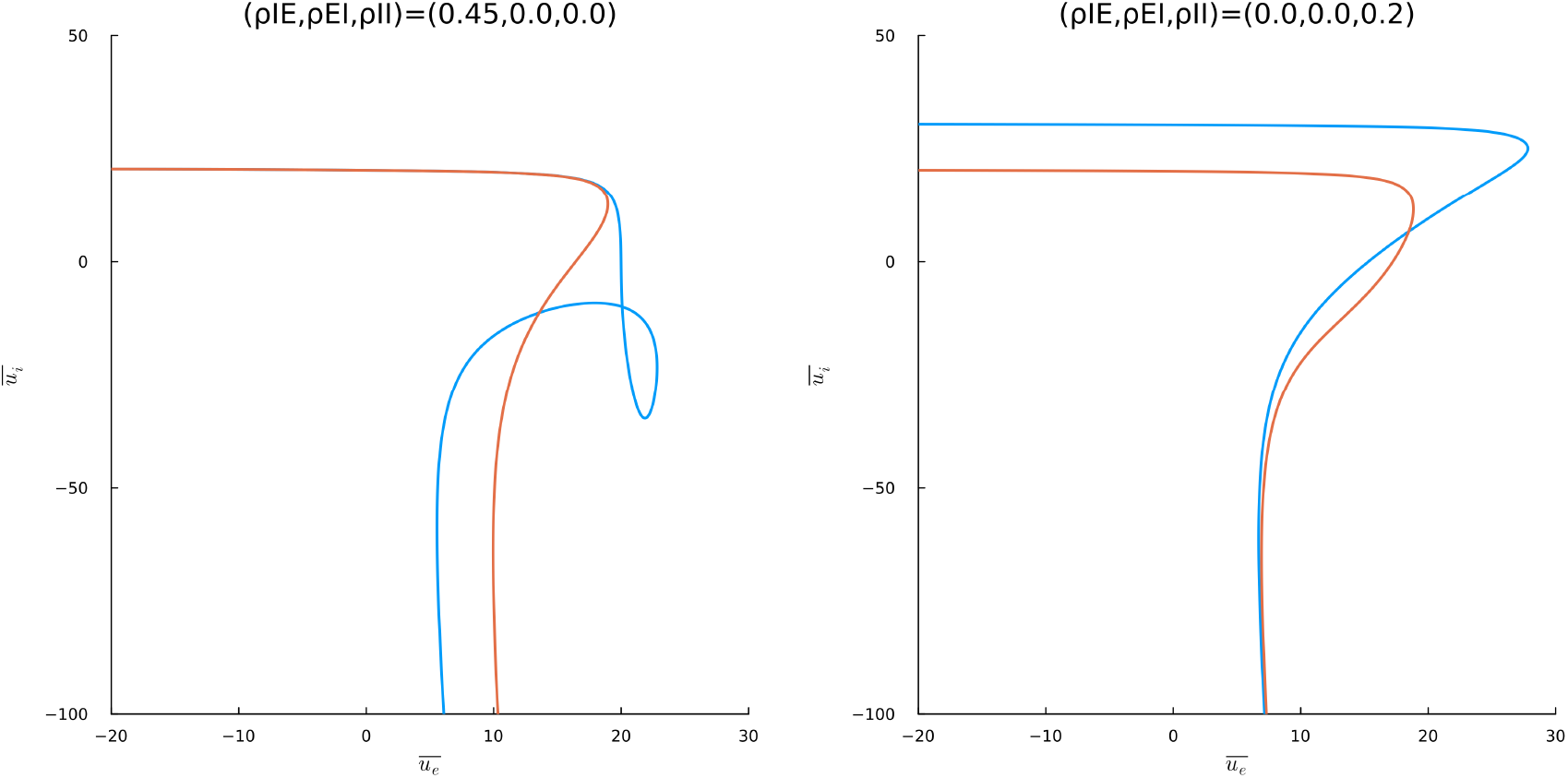
Input space Hopf bifurcation curves for **4POP WC-NM** with *γ* = 1. Here we have set *j*_*ee*_ = 0, *j*_*ei*_ = −40, *j*_*ie*_ = 40, *j*_*ii*_ = −40, Δ = 1, *τ* = 0.5, *D*_*i*_ = 0.2, *D*_*c*_ = 0.4. Red = transverse, Blue = longitudinal; left = **4POP-IE**, right = **4POP-II**. The **WC-NM** Hopf curves depicted here can be compared directly with those from **QIF-NM** in Figure 4 of the main text.

Before proceeding, we note that exactly the same procedure works in the **4POP WC-NM** case to translate the weight space transverse/longitudinal Hopf curves into transverse/longitudinal Hopf curves in the input space. Importantly, since the transformations are all diffeomorphisms on the prescribed domains, the number of double Hopf points is preserved by the translation from weight to input space (at least for sufficiently small cross-hemispheric coupling, in which case Hopf curve self-intersections will not occur). We show the result of carrying this out in Figure S4; note the close similarity with the leftmost **QIF-NM** Hopf diagrams shown for **4POP-IE** and **4POP-II** in Figure 4 of the main text. This demonstrates directly how weight space bifurcations beget input space bifurcations, and hence how studying the former ultimately can yield information about the latter, which are more directly relevant to the phenomenon of PING-ING alternation.

### Some additional numerical experiments

To conclude, we return to bifurcations in weight space. The analysis above reveals that, at least in the limit *D*_*i*_ → 0^+^, and at least for the threshold-linear **WC-NM** network, the main determinant of the parameter regimes for which AP-ING/IP-PING may coexist is the ratio *D*_*c*_/*D*_*i*_ of cross-hemispheric to local inhibitory delays. Does this remain true for *D*_*i*_ = *O*(1)? What happens for the **QIF-NM** model? To address these questions, we use the approach described above to compute double Hopf points numerically. Given a fixed parameter set ***p***, one obtains a sequence of *k* frequencies *ω*_*I*_ = *ω*_0_, …, *ω*_*k*_ = *ω*_*P*_ over which the Hopf branch in *Q* varies, as well as a sign *ε*_*k*_ = ±1 associated with each frequency, corresponding to whether the Hopf bifurcation bordering the stability region at that frequency is longitudinal/transverse, respectively. This technically provides correct information only in the weak coupling limit *ρ*_*xy*_ → 0^+^ but we have observed empirically that these signs remain accurate for *ρ*_*xy*_ positive but not too large (e.g. remaining bounded below 1/2). A natural measure of the similarity between the Hopf profile thus obtained, and the experimentally observed spectral profile, is

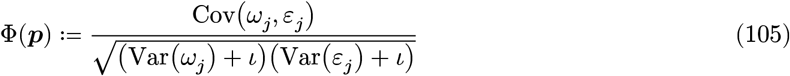

where Cov and Var denote sample covariance and sample variance of the vectors ***ω*** and ***ε***, while *ι* = 10^−8^ is a small parameter introduced for numerical stability. This is just a correlation between frequency and sign; **Φ** will be large and negative if oscillations at higher frequencies tend to be anti-phase and oscillations at lower frequencies tend to be in-phase, or large and positive if the opposite holds. Moreover, at parameter sets for which there are many double Hopf points, and correspondingly many in-phase/anti-phase transitions, these will cancel out and give a **Φ** value of lower magnitude. Hence, large negative values of **Φ** pick out precisely the portions of the parameter space most similar to the RSG profile.

**Figure S5:**
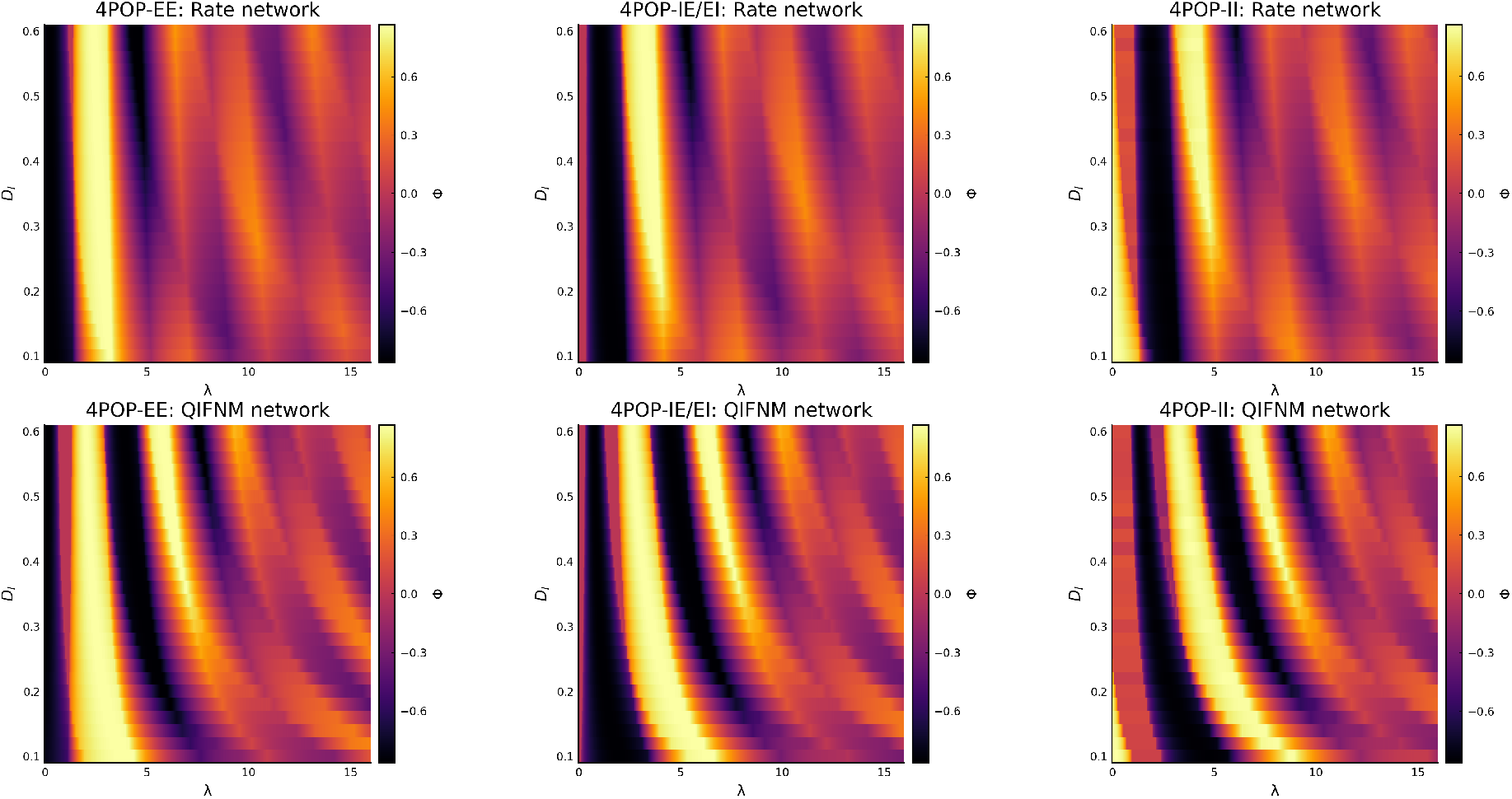
**Φ** computed for threshold-linear and QIF-NM **4POP** networks, varying *D*_*i*_ and *λ* = *D*_*c*_/*D*_*i*_. Here *τ* = 1, and for the QIF-NM we take *τ*_*m,E*_ = *τ*_*m,I*_ = 1, 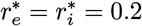. Large negative values of **Φ** correspond to parameter sets exhibiting high-frequency anti-phase and lower-frequency in-phase oscillations, similar to experimental data. Note the predictions from *D*_*i*_ → 0^+^ approximate well the *λ* ranges of large negative **Φ** for the rate network, while remaining qualitatively accurate for the QIF-NM networks.

In Figure S5, we show the numerically computed value of **Φ** across a wide range of *D*_*i*_ and *λ* pairs; regions with large negative **Φ** are in good approximate agreement with the asymptotic prediction from above. Remarkably, the results for the **QIF-NM** network are similar; in particular, the qualitative ordering of “RSG-like” delays supported by each model type (i.e., **4POP-EE** < **4POP-IE/EI** < **4POP-II**) is preserved.

As noted earlier, in the **4POP WC-NM** networks, the steady state rates, through the input gains *g*_Δ_, play a role which is essentially separable from that of synaptic kinetics. However, due to the rate-induced phase effects in **4POP QIF-NM** networks, the instabilities presented for the **QIF-NM** networks depend nontrivially on the values of the steady state rate parameters 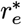 and 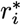. The values chosen in Figure S5 are 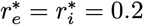, below the resonance threshold 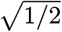. To examine the effect of population-level resonance on this double Hopf landscape, we fix the delays and vary 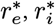, depicting the results in Figure S6.

**Figure S6:**
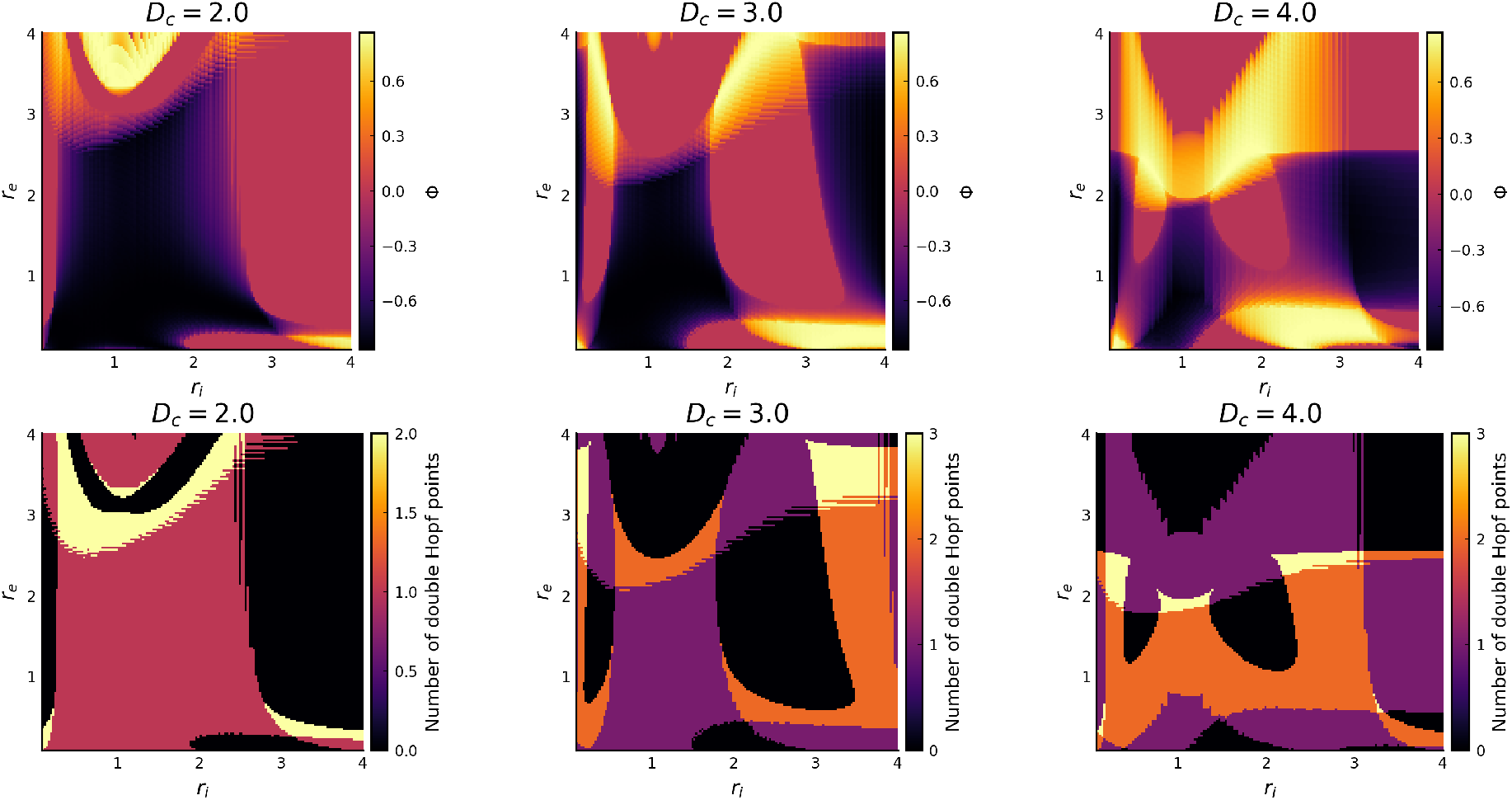
**Φ** computed for **4POP QIF-NM** networks, fixing delays and varying the resonance parameters *r*_*e*_, *r*_*i*_. Here we take *τ*_*m,e*_ = 9, *τ*_*m,e*_ = 5, *D*_*i*_ = 1 to match approximately the parameters from the main text. For short cross-hemispheric delays *D*_*c*_ = 2 − 3, we find a large region of the rate space presenting a single double Hopf point and large negative **Φ**. This region corresponds with that explored by varying the external inputs *η*_*E*_ and *η*_*I*_ in the main text, revealing the robustness of the preferred cross-hemispheric phase relationships to input variation. However, as the delay is increased, this region reduces in size, flanked by regions with “non-RSG-like” patterning, revealing a complex dependence on the resonant population frequencies.

**Figure S7:**
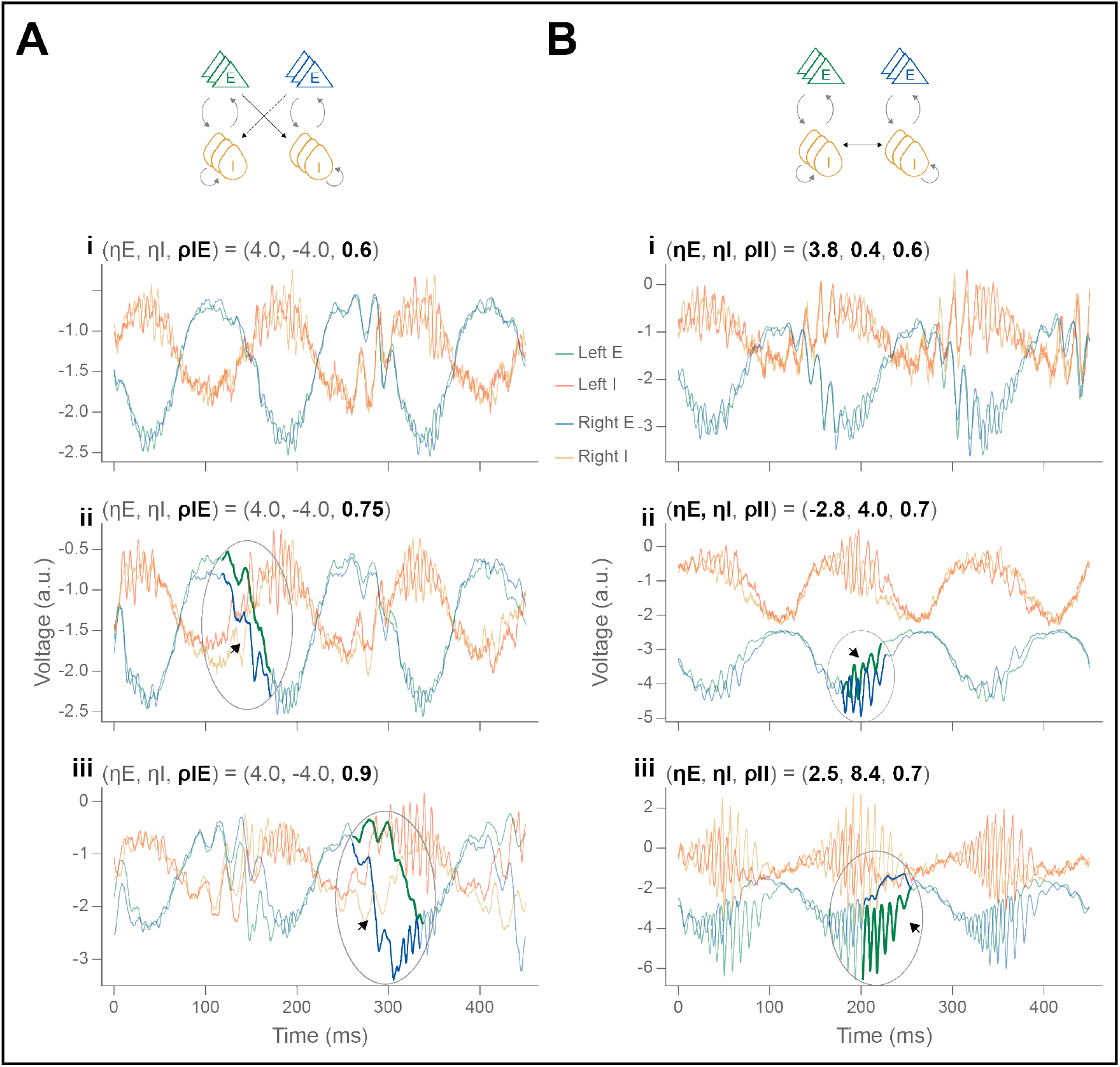
Dynamics of the mesoscopic model: IP-PING/AP-ING alternation and transient symmetry breaking. ***A***, simulation traces for **4POP-IE**, keeping (*η*_*E*_, *η*_*I*_ ) = (4, −4) fixed and varying *ρ*_*IE*_. **i**. At *ρ*_*IE*_ = 0.6, we observe robust IP-PING/AP-ING switching, with IP-PING and AP-ING amplitudes comparable for appropriate drive parameters (which does not seem achievable in the deterministic model, Figure 6D, ii). ii, At higher *ρ*_*IE*_ = 0.75, transient symmetry breaking events (TSB; pointed out by arrow and circle) appear, in which the mean voltage exhibits brief deviations across hemispheres. TSB events occur during the rising phase of the theta cycle, consistent with positioning of the transverse fold curve (Figure 5B, iii). **iii**. For even higher *ρ*_*IE*_ = 0.9, ING burst amplitudes may also differ across hemispheres, reflecting the ability for the unforced model to generate symmetry-broken ING (Figure 5B, iv, black marker). ***B***, simulation traces for **4POP-II. i**. an AP-ING/IP-PING switching solution in the noisy mesoscopic model. ii, For low *η*_*E*_, consistent with the positioning of the transverse fold curve for **4POP-II** (Figure 5C, iii), TSB occurs towards the end of the falling phase of the theta cycle. iii, At higher *η*_*E*_, TSB manifests as ING oscillations exhibiting marked amplitude and phase deviations during the falling phase of theta (akin to the out-of-phase ING oscillations in Figure 5C, iii, gold marker). Finite-size noise amplitudes in the mesoscopic model are given by *N*_*E*_ = *N*_*I*_ = 8000, while coupling strengths and interhemispheric delays agree with the default parameter values.

## Notes

### Competing Interest Statement

The authors have declared no competing interest.

